# Long-term B cell memory emerges at uniform relative rates in the human immune response

**DOI:** 10.1101/2023.11.27.568934

**Authors:** Ivana Cvijović, Michael Swift, Stephen R. Quake

**Author notes:** equal contribution (Ivana Cvijović, Michael Swift).

## Abstract

B cells generate pathogen-specific antibodies and play an essential role in providing adaptive protection against infection. Antibody genes are modified in evolutionary processes acting on the B cell populations within an individual. These populations proliferate, differentiate, and migrate to long-term niches in the body. However, the dynamics of these processes in the human immune system are primarily inferred from mouse studies. We addressed this gap by sequencing the antibody repertoire and transcriptomes from single B cells in four immune-rich tissues from six individuals. We find that B cells descended from the same pre-B cell (“lineages”) often co-localize within the same tissue, with the bone marrow harboring the largest excess of lineages without representation in other tissues. Within lineages, cells with different levels of somatic hypermutation are uniformly distributed among tissues and functional states. This suggests that the relative probabilities of localization and differentiation outcomes change negligibly during affinity maturation, and quantitatively agrees with a simple dynamical model of B cell differentiation. While lineages strongly co-localize, we find individual B cells nevertheless make independent differentiation decisions. Proliferative antibody secreting cells, however, deviate from these global patterns. These cells are often clonally expanded, their clones appear universally distributed among all sampled organs, and form lineages with an excess of cells of the same type. Collectively, our findings show the limits of peripheral blood monitoring of the immune repertoire, and provide a probabilistic model of the dynamics of antibody memory formation in humans.

---

The human antibody response provides protection against pathogens. The stability of this protective memory varies widely between pathogens, vaccines, and individuals, lasting for periods of times ranging from months to decades [1]. The durability of antibody memory is determined by the creation and persistence of the B cells that produce antibodies and is of great interest for vaccine development [2–4].

Human B cells exist in many different functional states or cell types, which have been characterized through immunophenotyping and single-cell sequencing studies [5–9]. These states differ in their ability to survive, secrete, and further evolve their antibodies in future immune responses. There are two major B cell memory states generated in immune responses: memory B cells and long-lived antibody secreting cells (ASCs). Memory B cells persist for long periods of time without secreting antibodies but can rapidly differentiate into ASCs when re-exposed to pathogens [10, 11]. In contrast, terminally differentiated long-lived ASCs constantly secrete antibodies regardless of whether there is an ongoing immune response [12, 13]. Unlike memory B cells, they are rarely found circulating in the peripheral blood and are thought to primarily reside in the bone marrow [14]. Both cell types are generated from Naive B cells in microanatomical structures known as germinal centers (GCs) that arise during an immune response [15].

Work in model organisms has described the process of affinity maturation and B cell differentiation in the GC in great detail [15]. Generally, B cells undergo rounds of random mutation of their antibody nucleotide sequences followed by competition on the basis of the affinity of their antibody to receive proliferation and differentiation signals from T cells. Differences in these signals direct them to become short or long-lived ASCs or memory B cells [15, 16]. A similar set of signals directs these cells to their particular niches, such as the bone marrow, after leaving GCs [14].

Less is known about the temporal and spatial dynamics of B cell memory formation in healthy humans. In mice, there remain two opposing paradigms for the pace of differentiation in the GC reaction [15]. Lines of research have provided support for a temporal switch in the GC reaction, in which memory B cells are favored early in the GC response and ASCs arise preferentially later [17, 18]. Other work has provided evidence that the generation of long-lived ASCs and their localization to the bone marrow is uniform in time during an immune response [19, 20]. It is not clear to what degree either of these paradigms applies to humans.

Antibody repertoire sequencing can provide a statistical view of human B cell evolution and differentiation. For example, repertoire sequencing has identified the genetic relationships between B cells in the blood [21], during vaccination [22–24], the order and pace of class-switch recombination [25], and the genetic relationships between B cells in different tissues [23, 26, 27]. However, insight into the dynamics of long-term memory formation has been limited because previous work has often sampled only the blood on short timescales (which explicitly omits long-lived ASCs), could not associate antibody sequences with cell states, or was not sufficiently powered for this type of analysis [6, 7, 28]. This gap leaves it impossible to systematically understand, for example, when long-lived ASCs arise during an immune response and how they are related to the B cells that circulate in the blood.

Here we use paired single-cell transcriptome and VDJ sequencing to measure the phenotypic and genetic relationships of B cells in four immune-rich tissues in six healthy individuals. We then use these measurements to investigate the repertoire-wide patterns of B cell evolution, differentiation, and migration between organs. We find that B cells make independent differentiation decisions at constant relative rates throughout the course of affinity maturation. Thus, long-lived ASCs are routinely generated at any point in immune responses. Furthermore, groups of related cells have highly correlated localization outcomes, meaning that tissue-resident cells are often found in large lineages that are not observed in the peripheral blood.

## The distribution of B cells in lymphatic tissues at rest

We purified B cells from the vertebral bodies, supradiaphragmatic and mesenteric lymph nodes, the spleen, and the blood of 6 organ donors with no clinical history of cancer, immune disease, or active infectious disease at the time of death (**Fig. 1**, **Table S1**, **Methods Summary**). We used droplet-based single cell sequencing to analyze the transcriptomes and VDJs of 200 thousand single B cells. To increase the depth of VDJ sampling, and to provide replication of the measurement process, we used the same platform to sequence only the VDJs of an additional 600 thousand B cells.

**Figure 1.**
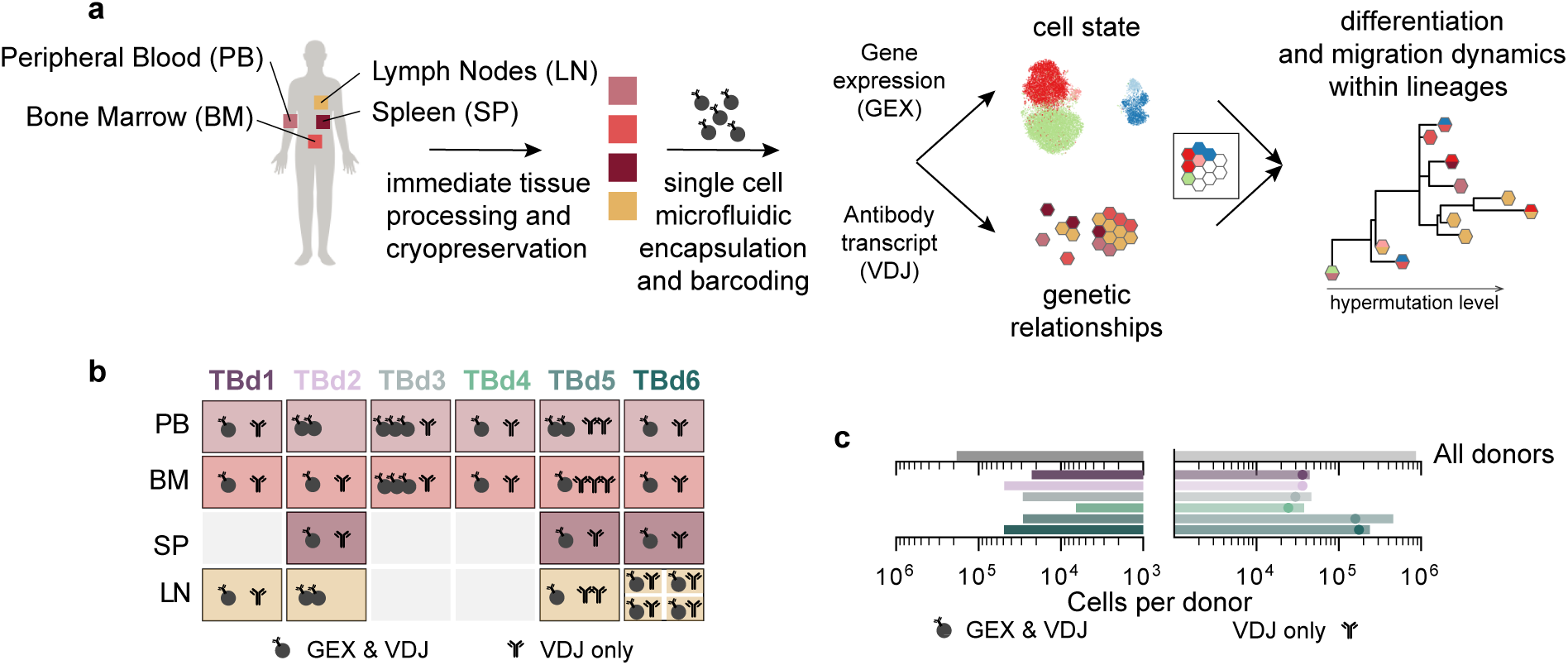
Overview of the study. (a) Study design. (b) Graphic depicting the data collected for each tissue and donor. GEX denotes single cell transcriptome sequencing and VDJ denotes antibody transcript (or, equivalently, the B cell receptor (VDJ)) sequencing. Note that for donor TBd6, we collected samples for four separate lymph nodes: three supradiaphragmatic lymph nodes, and one mesenteric lymph node. TBd2 and TBd5 lymph node samples were all derived from a single, individually processed supradiaphragmatic lymph node, whereas TBd1 lymph node samples represent the pooled mixture of three mesenteric lymph nodes. (c) Total numbers of cells in transcriptionally profiled samples (left) and single cell VDJs samples (right, see **Supplementary Information Sections 2 and 3** for a description of our cell calling algorithms). Dots represent the total number of unique VDJs in single cell VDJ only samples.

Overall, we observe few activated B cells in any donor, consistent with a quiescent immune system (**Extended Data Fig. ED1**). Activated B cells were also not more abundant in donor TBd1, who received a Moderna COVID-19 booster within 30 days prior to tissue recovery, consistent with ongoing affinity maturation in this donor remaining localized to the draining lymph nodes [29]. A small fraction of all B cells (1-6% per donor) appear to be proliferating as evidenced by the expression of M-phase markers (**Extended Data Fig. ED2**, **Supplementary Information Section 2.3**). Proliferating cells are predominantly of the ASC type, though we sampled a small number of proliferating GC B cells, as well as a small fraction of atypical B cells (ABCs) [9] that appear to be actively proliferating(**Extended Data Fig. ED2a**). The distribution of B cell subtypes among the sampled organs was consistent with previous findings [7]: the secondary lymphatic organs contained an abundance of memory B cells relative to other sampled organs, and the bone marrow was enriched for antibody-secreting cells (**Extended Data Fig. ED1**).

## The distribution of related B cells among lymphatic tissues

B cells descended from the same VDJ recombination event carry a unique genetic recombination signature that we used to group B cells into lineages (see **Supplementary Information Section 3.1**). We found that cells belonging to the same lineage tend to co-localize within the same tissue (see **Fig. 2a,b**). This suggests that related B cells receive consistent localization signals. In all donors except TBd1 (**Extended Data Fig. ED3**), this co-localization is particularly strong in the case of the bone marrow (**Fig. 2b**).

**Figure 2.**
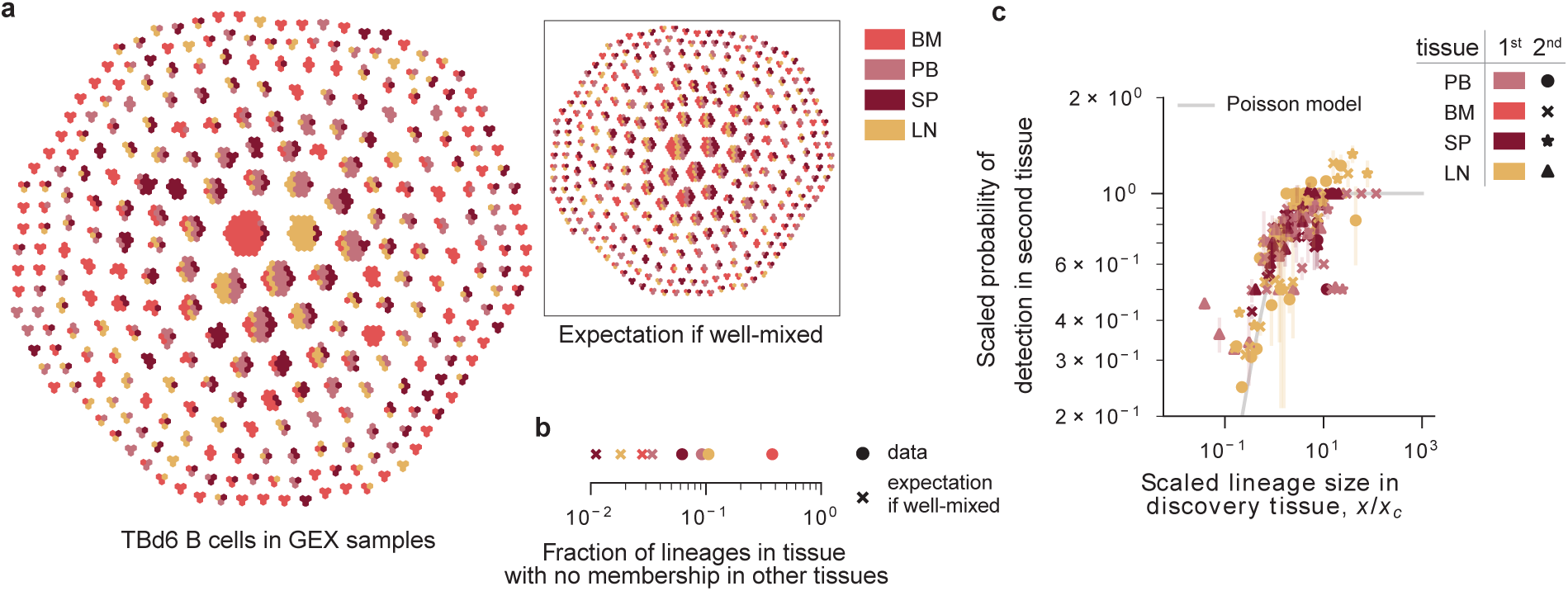
Tissue distribution of B cells descended from the same VDJ recombination event (lineages). (a) The tissue composition of lineages from TBd6. Each hexagon denotes a cell, and each connected group of hexagons represents a lineage. Cells are colored by tissue. Each tissue has been downsampled to the same overall number of cells. Inset shows the expectation if the lineages were well-mixed among the organs, obtained by randomly permuting the tissue labels among the cells. (b) The fraction of lineages in a tissue without sampled members in other tissues. Crosses denote well-mixed expectation, obtained by permutation. (c) The probability that a lineage is discovered in a tissue, as a function of the total size of that lineage in another, ascertainment, tissue. Points are colored by ascertainment tissue, and each panel corresponds to a different discovery tissue. Size represents the total fraction of unique heavy chain sequences attributable to the lineage, and is scaled by a single parameter, the critical size *x_c_*, shown in **Extended Data Fig. ED2**. As in Fig. 6, all independently generated emulsions have been downsampled to an equivalent number of unique VDJs per emulsion (5000), and error bars represent standard errors among the same quantity calculated using all available independent pairs of emulsions. Grey line denotes expectation if changes in localization are a Poisson process, 1 *− e^−x/x_c_^*.

However, B cell co-localization is not absolute: in large lineages, a minority of cells can be found in a different organ. This sharing follows a simple pattern: the probability that a lineage sampled in the blood, the spleen, or a lymph node, is present in a second tissue increases with its size (see **Fig. 2c**). The functional form of this increase suggests that the migration of B cells out of these tissues is governed by a Poisson process, where there is a small, but constant probability that a cell in a lineage found in one of these three tissues localizes in a different tissue. Once again, we note a departure from this overall pattern in TBd1. In this donor, we observe unusually large lineages and unusually high levels of lineage sharing between the bone marrow and the blood (**Extended Data Fig. ED3**), perhaps related to a transient process associated with their recent COVID-19 booster.

## Difference in hypermutation levels among related cells in different tissues

Overall, these results suggest that B cells belonging to the same lineage get consistent localization signals at the point of GC exit, but that a portion of the lineage can still localize in a different tissue. We wanted to understand whether there is covariation between hypermutation and tissue localization (**Fig. 3**). Such covariation could, for example, be mediated by antibody affinity for antigen, which both increases with hypermutation, and affects cell differentiation potential: cells with the highest affinity are thought become ASCs, a cell type that often localizes to the bone marrow [14, 15].

**Figure 3.**
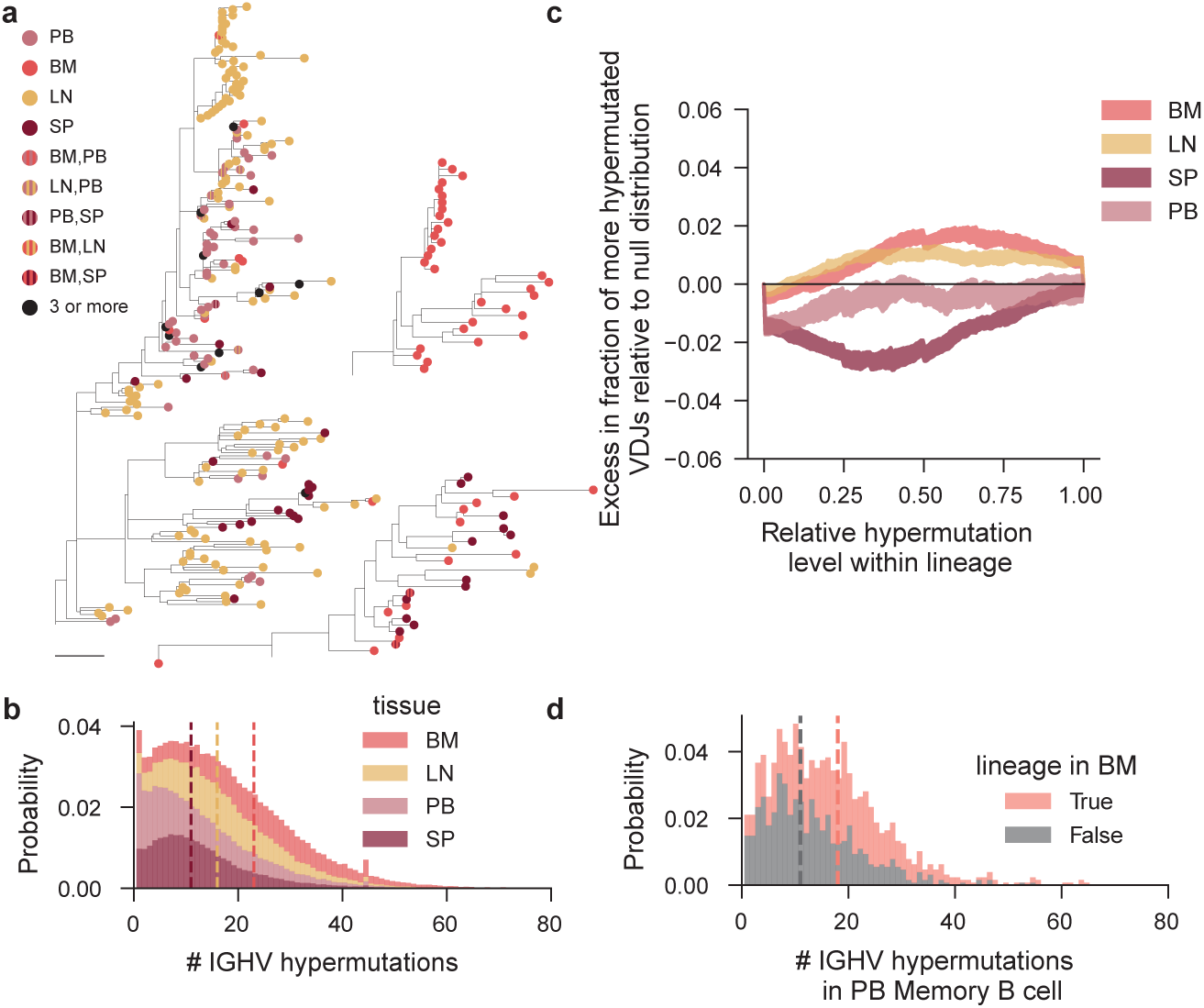
Distribution of hypermutation levels among related cells found in different tissues. (a) Templated V gene trees for 3 example lineages from TBd6. Trees are rooted on the inferred germline V sequence. Leaves correspond to unique templated V sequences and are colored by localization. Bar denotes a divergence of 5%. (b) Distribution of the number of hypermutations observed in the IGHV gene for all unique heavy chains, split by tissue. Dotted lines represent medians, which overlap for blood and spleen in our data. (c) The within-lineage distribution of hypermutated cells in a particular tissue. Lineages with at least 3 unique VDJs are included in this analysis. For each unique VDJ, we quantified its relative hypermutation level within the lineage: 0 and 1 correspond to the least and most hypermutated sequences in the lineage, respectively. We obtained null distributions of this quantity by permuting tissue labels between VDJs associated with lineages containing that tissue (e.g. the BM labels were permuted among lineages with at least one member sampled in the bone marrow). The confidence intervals represent the bounds on the cumulative density function obtained in 100 independent permutations. We plot the difference between the measured and null distributions. Black denotes perfect agreement. (d) Distribution of the number of hypermutations observed in the templated portion of the IGHV gene for VDJs found to be associated in peripheral blood memory B cells. Grey and salmon represent memory B cells with and without a sampled relative in the bone marrow, respectively.

Notably, the overall level of hypermutation does vary between cells residing in different tissues, and B cells located in bone marrow were, by far, the most hypermutated (**Fig. 3b**). Surprisingly, within lineages, cells sampled in different tissues were almost entirely uniformly distributed in hypermutation level (**Fig. 3c**). Specifically, bone marrow localized cells could appear at any point in the lineage, and were only weakly enriched among the most strongly hypermutated cells – the percentage of bone marrow cells that have a relative hypermutation level greater than 0.5 deviates from the null distribution by only on the order of 1%. This apparent incongruity is explained by the observation that hypermutation levels co-vary between B cells in the same lineage. For instance, if a lineage is detected in the bone marrow, members of that lineage outside of the bone marrow are more hypermutated than comparable cells. (**Fig. 3d**).

The results in **Fig. 3c** suggest that decisions to localize in a certain tissue change only weakly during affinity maturation. However, tissues are mixtures of B cells of different cell types. Thus, conditioning on tissue presence of a lineage still recovers a mixture of lineages that have experienced different amounts of affinity maturation: both naive lineages, and very mature ones, in the case of the bone marrow. Thus, it is hard to extract meaningful dynamical insight without knowledge of the underlying cell states. In the next section, we explore the underlying differentiation process of the individual B cell subtypes, which allows us to describe the dynamics of memory formation.

## Phenotypic variability among non-naive B cells

We initially labeled the cell types using `celltypist`, an algorithm for automatically assigning cell types to gene expression profiles [7]. Our data revealed additional phenotypic variability within ASCs (ASCs, see **Extended Data Fig. ED4** and **Extended Data Fig. ED5**), consistent with a prevailing understanding that ASCs harbor substantial functional diversity [5, 14, 30]. We detected ASCs expressing all isotypes. Interestingly, we also observed a fraction of IGHD+ ASCs (**Extended Data Fig. ED4d**); these ASCs were more heavily hypermutated than ASCs of other isotypes, and likely represent a class-switched IgD population [31]. ASCs formed four distinct clusters (ASC-1 through ASC-4) with notable differences in their gene expression (**Extended Data Fig. ED4a,b**). A substantial fraction of ASC-3 were proliferating and had a gene expression signature that closely resembles previous descriptions of plasmablasts [32] (see **Extended Data Fig. ED4b**). The other types of ASCs appeared non-proliferative and were commonly found outside the peripheral blood (**Extended Data Fig. ED4b,c**). In particular, ASC-1 are primarily located in the bone marrow and have the canonical markers of long-lived plasma cells [5, 33](**Extended Data Fig. ED4b,c**).

## Uniform differentiation rates during the GC reaction

Similarly to the observations we made with respect to B cells located in different tissues, we find systematic differences in the hypermutation levels of different B cell types (**Fig. 4a**). However, within lineages, we cannot detect statistically significant differences in the level of hypermutation between related cells of different types, with the exception of Naive B cells (**Fig. 4b**, **Extended Data Fig. ED6a**, note the same does not apply to B cells of different isotypes, see **Extended Data Fig. ED6b**). This surprising observation suggests that, during affinity maturation, the relative probabilities of differentiation into a memory B cell or an ASC remain consistent. In other words, they argue against a “temporal switch model” in which ASCs arise preferentially later in the course of affinity maturation [17].

**Figure 4.**
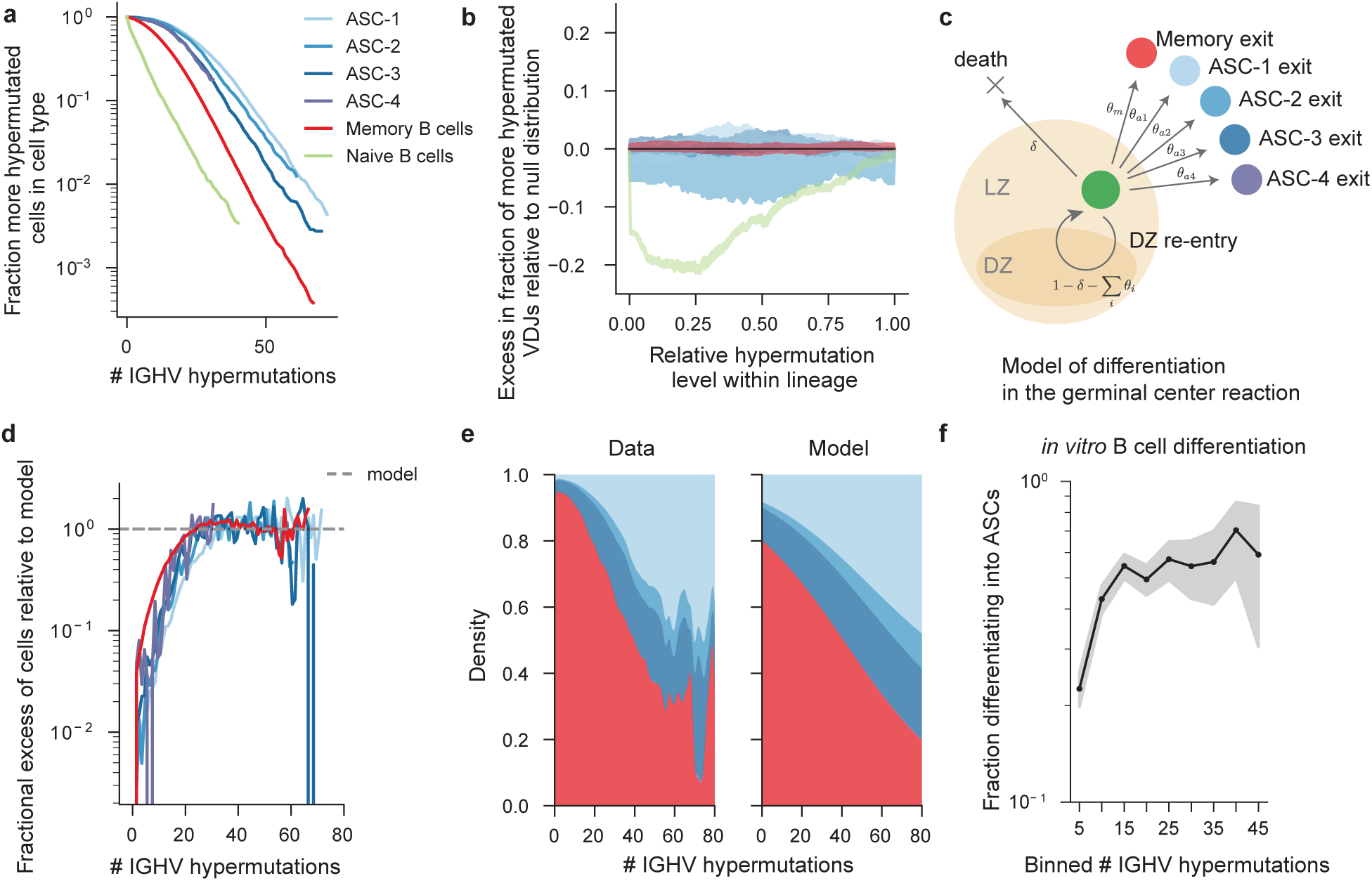
Distribution of hypermutation levels among related cells of different cell types. (a) Distribution of the number of IGHV hypermutations for heavy chains associated with different cell types. Cells with no observed hypermutations are excluded, for consistency with the following panels. (b) The within-lineage distribution of hypermutated cells in a particular cell type, as in Fig. 3. (c) Schematic of a simple model of differentiation in the GC reaction in which GC B cell make cell fate decisions at rates that do not change as the GC reaction proceeds. For each cell fate outcome, the per-cycle rates are denoted in symbols adjacent to the arrows. (d) Quantitative agreement between measured hypermutation distribution and constant exit rate model with the best-fit parameters. (e) Fraction of cells of each cell type as a function of the hypermutation level. Left panel represents the empirical distribution, and the right represents the distribution expected assuming the model in (c) with the best-fit exit rates and matched overall abundances of B cells of each type. Colors as in previous panels. (f) Differentiation statistics of human B cells *in vitro*: probability of differentiating into an ASC across hypermutation levels re-analyzing *in vitro* data from [34]. Shaded areas denote the 95% confidence intervals and are binomial proportion intervals calculated using the Clopper-Pearson method and bins containing fewer than 10 cells are omitted.

To investigate this hypothesis further, we formulated the simplest possible model of differentiation during the GC reaction consistent with these observations. Specifically, we considered a model in which B cells in the GC have a constant probability of dying or exiting the GC as one of the B cell subtypes every time it re-enters the light zone of the GC (i.e. upon the completion of each “GC cycle”, see **Fig. 4b**). In this model, the distribution of hypermutation levels for each of the B cell subtypes is expected to have an exponential tail with a rate that is proportional to the per-cycle exit probability (see **Supplementary Information Section 6** for details). This prediction agrees with the measured hypermutation distributions over several orders of magnitude, for cells that have accumulated over 10 hypermutations (**Fig. 4d, Supplementary Figure S25**). At lower hypermutation levels, all cell types deviate from this “constant exit rate” model in a consistent way, which suggests there is an overall change in the rate at which long-term B cell memory is formed in the early GC. However, because the reduction in the exit rate is consistent for the different cell types (**Fig. 4d**), the relative rates at which memory B cells and ASCs are created remains remarkably consistent throughout the GC reaction, with memory B cells being created in each GC cycle at an estimated rate that is roughly 40% higher than that of ASCs (see Table S3 in the **Supplementary Information**).

This difference in relative exit rates can explain the observation that ASCs tend to be more hypermutated than memory B cells: since differentiation into an ASCs occurs with a lower probability than differentiation into memory B cells, we expect ASC creation to occur after on average more GC cycles (see **Fig. 4e**, **Supplementary Information Section 6**). Finally, we compared these observations to the results of a previously published *in vitro* human B cell differentiation experiment [34]. In that work, we observed that hypermutated B cells were intrinsically more likely to differentiate into ASCs than unhypermutated B cells when stimulated with the same T cell-dependent stimulus *in vitro*. We re-analyzed these data to quantify how the differentiation bias changes across hypermutation levels, and found that the probability of differentiating into an ASC is uniform across hypermutation levels, if a B cell has accumulated order 10 hypermutations prior to *in vitro* differentiation (see **Fig. 4f**). This experimental observation shows that no population-level changes in these intrinsic biases are observable *in vitro* across a range of hypermutation levels consistent with our observational data from human donors.

## Distributions of cell types within lineages

This view of differentiation in the GC implies that the cell type composition of a lineage can be predicted by the amount of hypermutation observed in the lineage. To verify this, we constructed a null probability distribution of B cell types as a function of the number of hypermutations observed in the templated IGHV gene from the data (**Fig. 4e**, left). We found that this model predicts both the probability that a lineage contains a cell in a certain state (see **Fig. 5a**), as well as the total number of cells found in a certain state as a function of lineage size (see **Fig. 5b**). Notably, this is not true for ASC-3s, which are found in far larger numbers in large lineages than expected. Thus, ASC-3s arise in a correlated fashion within certain lineages. In other words, there appears to be a global property of certain B cell lineages that may make them more prone to generating more ASC-3s.

**Figure 5.**
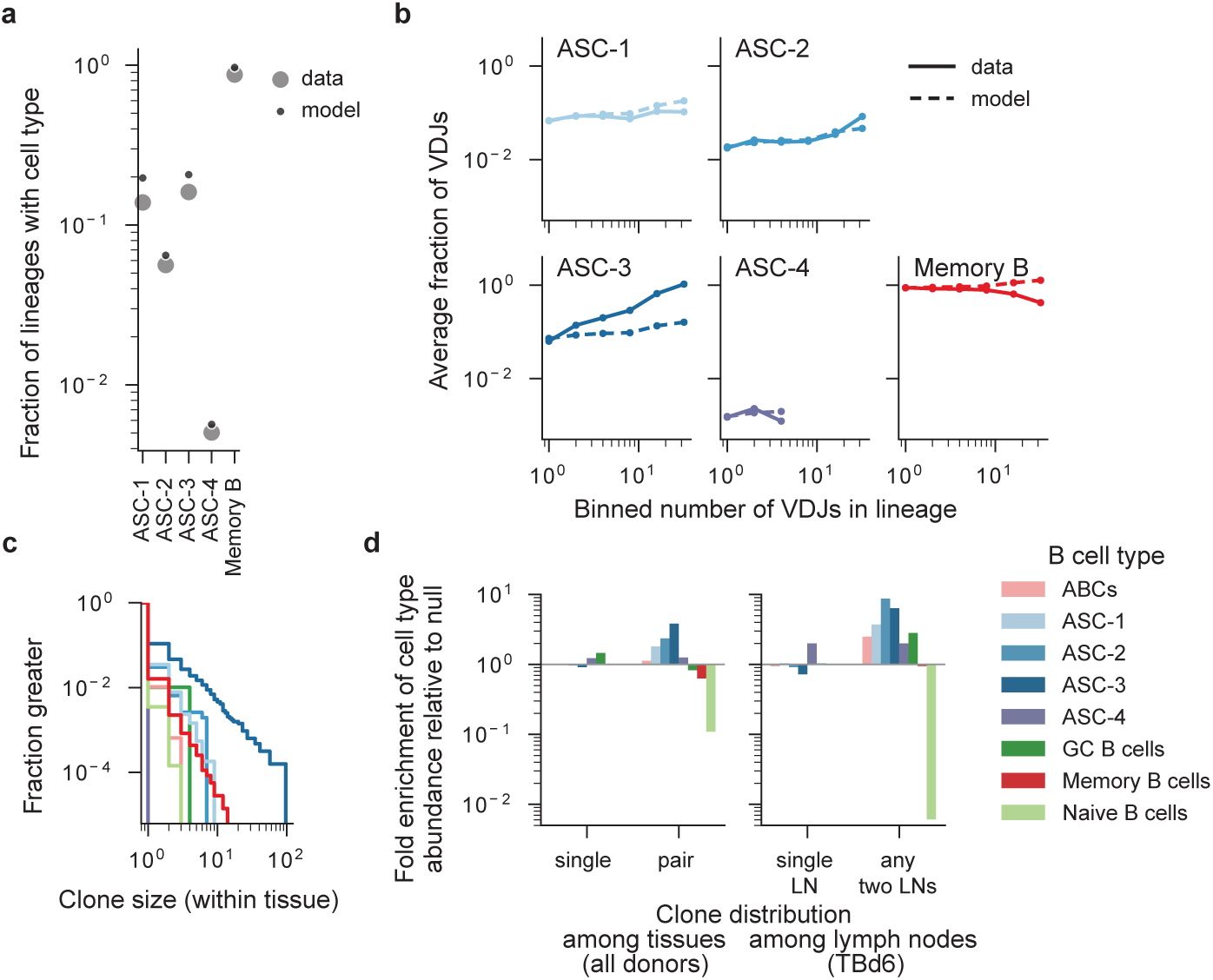
Distributions of cell types among cells belonging to the same lineage. (a) The total fraction of lineages with at least 2 unique VDJs that contained at least one cell of each type and (b) The average fraction of cells of a given type among lineages of a certain size. To construct the null expectations, we conditioned on the sampled phylogenies of all hypermutated cells with available transcriptomes and assumed independence between sampled cells. Points supported by fewer than 6 cells and fewer than 10 lineages have been omitted. (c) Clone size distributions of different B cell subsets. The clone size is the number of cells of each cell type that have an identical heavy chain nucleotide (VDJ) sequence and are found in the same tissue. Colors as in (d), which shows the changes in abundances of B cell subtypes as a function of the tissues their VDJ sequences were found in. As explained in **Supplementary Information Section 5.2**, VDJs are associated with a subtype label if they are found in a B cell of that subtype in any tissue. The null expectation is computed from the marginal subtype distributions across the tissues.

Given the resemblance of ASC-3s to plasmablasts, which are thought to arise during active immune responses, we wanted to understand whether ASC-3s preferentially arise in recently generated, ‘young’ lineages of B cells. To test this, we reasoned that, because human Naive B cells are thought to have a half-life only several weeks [35], lineages in which we observe an unhypermutated Naive B cell must be younger on average. Memory B cells sampled in these lineages typically carried fewer hypermutations (**Extended Data Fig. ED7a**), consistent with the interpretation that they are newly generated. However, not only did these “young” lineages not contain an abundance of ASC-3s, but ASC-3s accounted for a far higher proportion of cells in lineages that did not contain Naive B cells (**Extended Data Fig. ED7b**). This suggests that a global property other than age controls the abundance of ASC-3s in lineages. These findings are intriguing in light of a recent finding by Phad *et al.* [36], which showed that proliferative ASCs in the peripheral blood are often members of persistent lineages and are observable long after antigen encounter. Their observations likely pertain to the same ASC subtype, as ASC-3s account for almost all of the measured ASCs in the peripheral blood (see **Extended Data Fig. ED4**).

## Sharing of a subset of expanded ASC clones

Finally, we noticed that about 1 in 100 of all unique nucleotide VDJ sequences were identical in many individual B cells within the same donor, suggestive of extensive clonal expansion without hypermutation (**Extended Data Fig. ED9a**). This observation cannot be explained by the repeated generation of ‘public’ B cell clones, which arise at rates that are several orders of magnitude lower (**Supplementary Information Section 4**, **Extended Data Fig. ED8**). Though evidence of clonal expansion can be found among cells of all types, it is particularly prominent among ASCs, especially ASC-3s (**Fig. 5c**).

Our analysis of the tissue localization statistics of lineages showed that, when lineages reach a threshold size, they have an order one probability of being found in multiple tissues. We wondered whether expanded clones had a similar pattern. We quantified how often expanded clones were restricted to a single tissue. Once we accounted for the variability in the number of VDJs detected in each sample, we found that, across donors, all pairs of tissues shared similar fractions of VDJs (**Fig. 6a**, **Extended Data Fig. ED9b**). Since even replicates of the same tissue have limited overlap in the context of limited sampling, we derived a normalized probability estimate. This quantity constructs an empirical estimate of the sampling probability of an expanded VDJ using independent samples of the same tissue (see **Supplementary Information Section 5**). Surprisingly, VDJs expanded in one tissue are found in other tissues with normalized probabilities often exceeding 0.1 (**Fig. 6b**, **Extended Data Fig. ED9c**), and VDJs found in at least two tissues appear in a third, or fourth tissue with normalized probabilities of order 1. These observations suggest that expanded clones often achieve universal distribution throughout the sampled organs.

**Figure 6.**
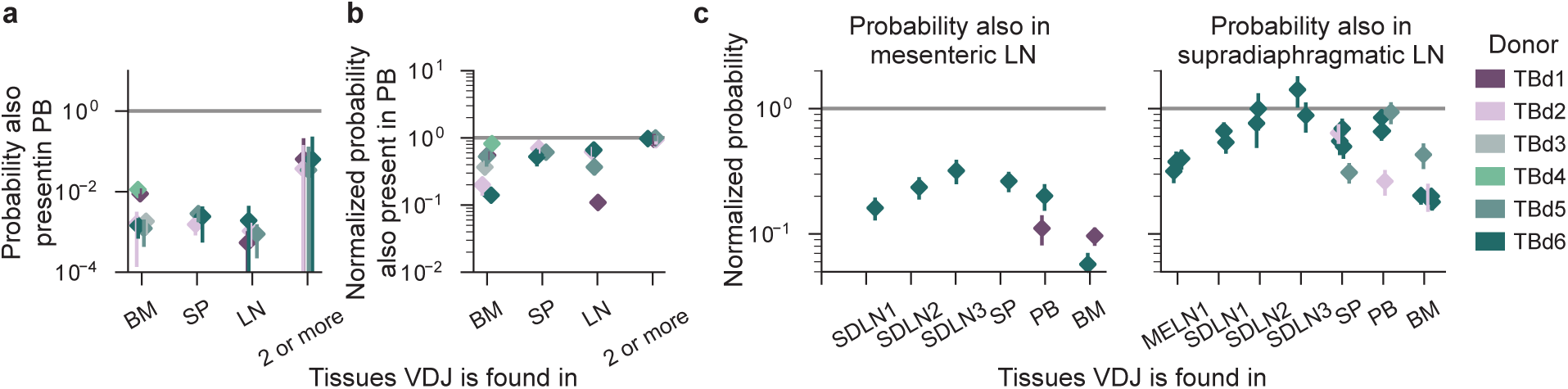
Localization and identities of shared B cell clones. (a) Probability that a B cell clone is found in the peripheral blood, given its VDJ has been sampled in a different tissue. In (b), we normalize the probability of appearance in the blood by the probability that the same VDJ was encountered in an independent sample of the tissue indicated on the *x*-axis (see **Supplementary Information Section 5** for details). This effectively conditions on expansion in the original tissue. To make symmetric comparisons between tissues sampled at different depths, all samples were downsampled to 3000 unique VDJs per independently generated 10X emulsion from a cell suspension, and smaller samples were dropped from the analysis. Individual points represent averages of 100 independent downsamples of the entire dataset, and error bars denote the standard error obtained from independent 10X runs of the same tissue, when available. (c) Probability that a B cell clone is found in a specific lymph node, conditional on it being expanded in another lymph node or non-lymph node tissue. MELN – mesenteric lymph node; SDLN – supradiaphragmatic lymph node

Earlier work revealed distinct networks of B cells associated with different anatomical locations [26], and we were interested if there was additional observable structure amongst lymph nodes from different anatomical locations. Consistent with these earlier findings, mesenteric lymph nodes shared notably fewer clones with all other tissues than supradiaphragmatic lymph nodes (**Fig. 6c**, **Extended Data Fig. ED9d**), and distinct supradiaphragmatic lymph nodes shared clones amongst themselves as often as they did with other tissues (**Fig. 6c**, **Extended Data Fig. ED9c**).

Multi-tissue clones are predominantly ASCs, and in particular ASC-3s (**Fig. 5d**). Still, we detected many shared memory B cell clones between tissues. We wondered if they also had distinctive gene expression features compared to memory B cells which were not shared between tissues. We found shared memory B cell clones had a gene expression program distinguished by MAST4 and a non-switched phenotype (**Extended Data Fig. ED10a,b**, **Supplementary Information Section 5.3**). Interestingly, the gene expression signature we identified using the Spleen and Lymph Node seemed to universally enrich for shared memory B cell gene expression amongst any pair of tissues containing a secondary lymphoid organ (**Extended Data Fig. ED10b,c**). **Discussion** In this work, we measured the VDJs and transcriptomes of B cells sampled from a number of organs from six human donors. This allowed us to investigate the statistics of B cell evolution, migration, and differentiation in the B cell system. We assess the amount of overlap in antibodies produced by B cells residing in different organs and find that related B cells tend to co-localize in the same lymphatic tissue. This effect was particularly prominent in the bone marrow, which contained the greatest enrichment of large, private B cell lineages not observed in the other lymphatic organs. Thus, there is substantial spatial structure in the B cell immune system that should inform the interpretation of peripheral blood draws for the purpose of B cell repertoire monitoring.

The co-localization of related B cells suggests that many of them acquire coherent localization signals during their generation. This coheres with observations made by [37] showing that qualities of B cell memory recall depend on the relative anatomical locations of the prime and boost during vaccination. However, a subset of ASCs (here labeled ASC-3) with properties similar to plasmablasts depart from this pattern. These cells collectively account for about 1 in 100 of all of the sampled VDJs and achieve universal distribution among the sampled organs. They represent the predominant mode of clone sharing between all sampled sites, including pairs of anatomically similar lymph nodes.

Though we have not sampled any lymph nodes with evidence of ongoing affinity maturation, we find that anatomically similar lymph nodes share memory B cell clones at rates no higher than other pairs of tissues. These findings have interesting implications for GC dynamics, especially in light of work showing that antibody evolution is shaped by secreted antibodies [38–40]. They suggest the evolutionary outcomes of GCs may be modulated through the effect of sharing of recently created, cycling ASCs from distant areas of the body.

Our analysis of long-term B cell memory shows that related B cells that have undergone different amounts of affinity maturation (as evidenced by their hypermutation levels) have remarkably uniformly distributed probabilities of localization and differentiation. It is possible that other, transient populations of differentiated B cells are created during immune response and deviate from these patterns, and studying their dynamics in light of our results is an interesting avenue for future work. In contrast with previous work that has suggested a temporal switch in which GCs become biased towards ASC production in the later phases of affinity maturation [17], our data are consistent with a simple model of B cell differentiation during affinity maturation in which differentiated subtypes of cells arise at a constant relative rate at all but the weakest hypermutation levels (*>* 3%). Moreover, more than 80% of differentiated B cells exit the GC at a point in affinity maturation when differentiation rates have stabilized (see **Fig. S15** in the **Supplementary Information**). Our observations are consistent with a recent study conducted in mice, which shows that long-lived ASCs continuously populate the bone marrow during an immune response [19, 41]. Thus, we suggest long-lived ASCs are created early during the human immune response as well, albeit more rarely than comparable memory B cells.

This observation raises interesting questions in light of the fact that antibody affinity is thought to be a primary determinant of B cell differentiation outcomes [15]. It is possible either that high-affinity antibodies can arise very early during affinity maturation, or that the affinity threshold governing differentiation potentials becomes more stringent as the GC reaction proceeds. These two pictures have different implications for both our understanding of the dynamics of affinity maturation as well as practical applications, including *in vitro* antibody design and vaccine design. Future work that combines single cell antibody repertoire sequencing with systematic measurement of the binding affinities along lineages would distinguish between these two pictures.

## Materials and methods summary

Fresh, whole, and non-transplantable organs were obtained from surgery and transported on ice by courier. Upon receipt, tissues were dissociated into single cell suspensions using methods developed by the Tabula Sapiens Consortium [28]. Some cell suspensions were processed immediately (fresh), and most were frozen and stored in liquid nitrogen. With the exception of the lymph nodes, all samples were enriched for the B cell lineage by selective depletion of non-B lineages by immunodepletion with magnetic beads. We used 10X 5’ immune profiling to encapsulate cells and barcode mRNA for single cell library prep. Cells were loaded at a target of 20,000 cells per lane for gene expression samples and 80,000 cells per lane for VDJ-only profiling. We used `cellranger 7.0.1` to generate cell by gene count tables and VDJ assemblies followed by custom pipelines to process the gene expression data and assemblies. All cell types were annotated manually using input from both the gene expression data and the VDJ sequencing data. To ensure reproducibility, we used the Snakemake workflow manager [42]. Detailed descriptions of all experimental protocols and analyses are provided in the **Supplementary Information**. The code we developed will be publicly available on Github at time of publication.

## Acknowledgments

We express our gratitude to all of the anonymous organ and tissue donors and their families, for giving the gift of life and the gift of knowledge, by their generous donation. We thank Donor Network West for their partnership on this project, and Bob Jones and Sheela Crasta for assistance with donor coordination and the logistics of tissue procurement. We are grateful to Zachary Sethna for help in implementing OLGA and estimating the selection factor used in interpreting the amount of VDJ sharing across donors. We thank Daniel Fisher, Ben Good, Felix Horns, Elizabeth Jerison, Mira Moufarrej, Alex Nguyen Ba, Alana Papula, John McEnany, and Soso Xue for useful discussions and helpful comments on the manuscript. This project has been made possible in part by grant nos. 2019-203354, 2020-224249, and 2021-237288 from the Chan Zuckerberg Initiative DAF; an advised fund of Silicon Valley Community Foundation; and by support from the Chan Zuckerberg Biohub. MS was supported by the NSF Graduate Research Fellowship. IC was supported by the Stanford Science Fellowship. Some of the computing for this project was performed on the Sherlock cluster. We would like to thank Stanford University and the Stanford Research Computing Center for providing computational resources and support that contributed to these research results.

## Author Contributions

IC & MS: Conceptualization, Methodology, Software, Formal Analysis, Investigation, Writing – original draft, Writing – review & editing. SRQ: Supervision, Funding Acquisition, Writing - review & editing.

## Data and code availability

Raw sequencing reads will be deposited in the NCBI BioProject database upon publication. All associated metadata, as well as the source code for the raw data processing pipeline, downstream analyses, and figure generation, will be made available at GitHub upon acceptance. Processed AnnData objects will be available on the CellxGene Portal pending acceptance.

---

**Table S1.** Demographic characteristics of donors.

| Donor | Age | Sex | Ethnicity | Cause of Death |
| --- | --- | --- | --- | --- |
| TBd1 | 46 | F | Asian/Japanese | Cerebrovascular stroke |
| TBd2 | 35 | M | American Indian or Alaska Native | Anoxia |
| TBd3 | 25 | F | Hispanic/Latino | Head Trauma |
| TBd4 | 43 | F | Hispanic/Latino | Anoxia |
| TBd5 | 45 | F | Asian / Filipino | Cerebrovascular stroke |
| TBd6 | 60 | M | Asian / Filipino | Anoxia, spontaneous cardiac arrest |

**Figure ED1.**
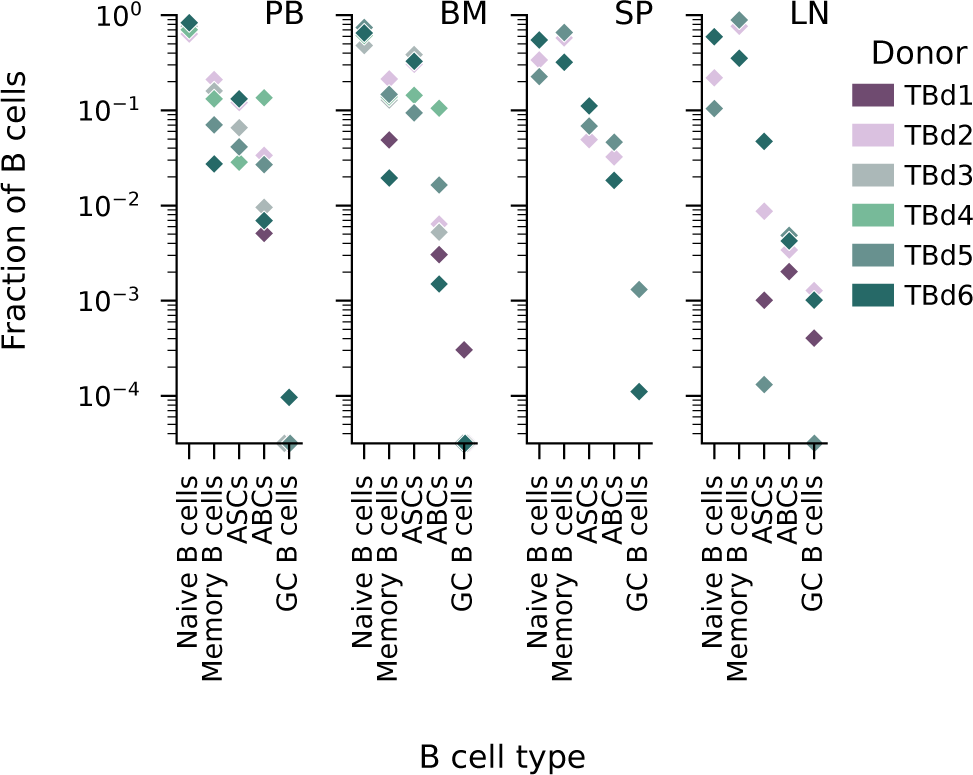
B cell type distribution across donors and tissues. Displayed are cell states represented by more than 5 cells across all donors. Pre and pro B cells are discussed in **Supplementary Information Section 2.6**. In cases in which a cell type in a sampled tissue was not detected among the sampled B cells, the symbol for that donor is shown on the x-axis. ASC - antibody secreting cell, ABC - atypical B cell, GC B cell - germinal center B cell.

**Figure ED2.**
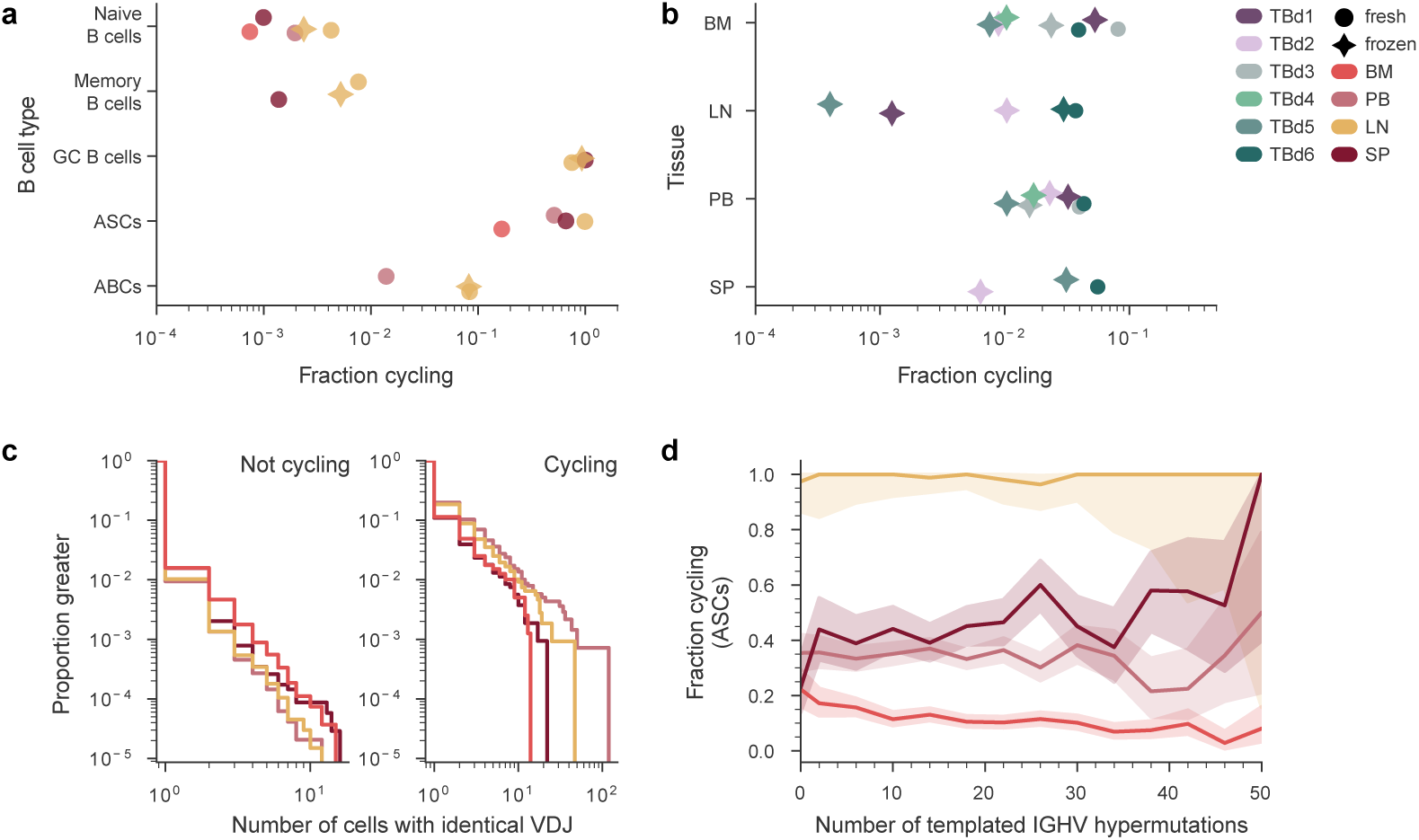
Cell cycle analysis across tissues. (a,b) Fraction of cells of each cell type and tissue (a) and tissue and donor (b) that are cycling. Markers denote whether a particular sample of a tissue was prepared fresh or previously cryopreserved. (c) The distribution of numbers of cells with identical VDJ sequences plotted by whether any or none of the sampled cells is found to be cycling. (d) Fraction of ASCs that are cycling shown for different tissues across hypermutation levels. Shaded areas denote the 95% confidence intervals are binomial proportion intervals calculated using the Clopper-Pearson method. Colors as in (b)

**Figure ED3.**
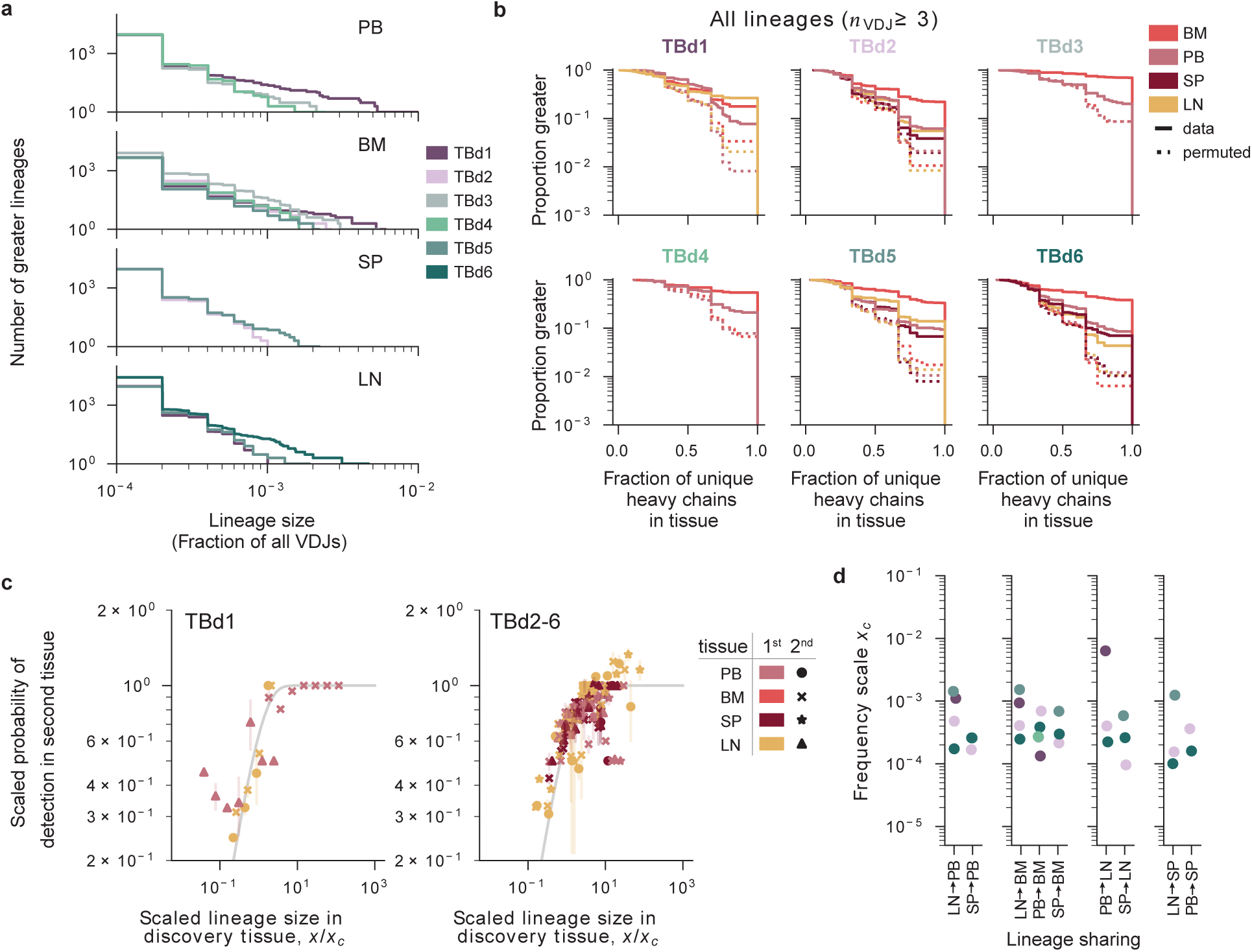
Distributions of cell types among cells belonging to the same lineage. (a) Lineage size distributions for the data used in **Fig. 2(c)** (b) The distribution of the fraction of unique VDJs in the lineage found in a certain tissue. Each panel is a different donor. Dotted lines denote the distributions obtained after a random permutation of the tissue labels within the same donor among the sampled cells. Each tissue has been down-sampled to the same overall number of cells. (c) The scaled probability that a lineage is discovered in a tissue, with anomalous donor TBd1 separated. Data plotted as in **Fig. 2(c)**. (d) Inferred frequency scale *x_c_* from **Fig. 2(c)** for each between pairs of organs where point are estimates for each donor (colors as in (a)).

**Figure ED4.**
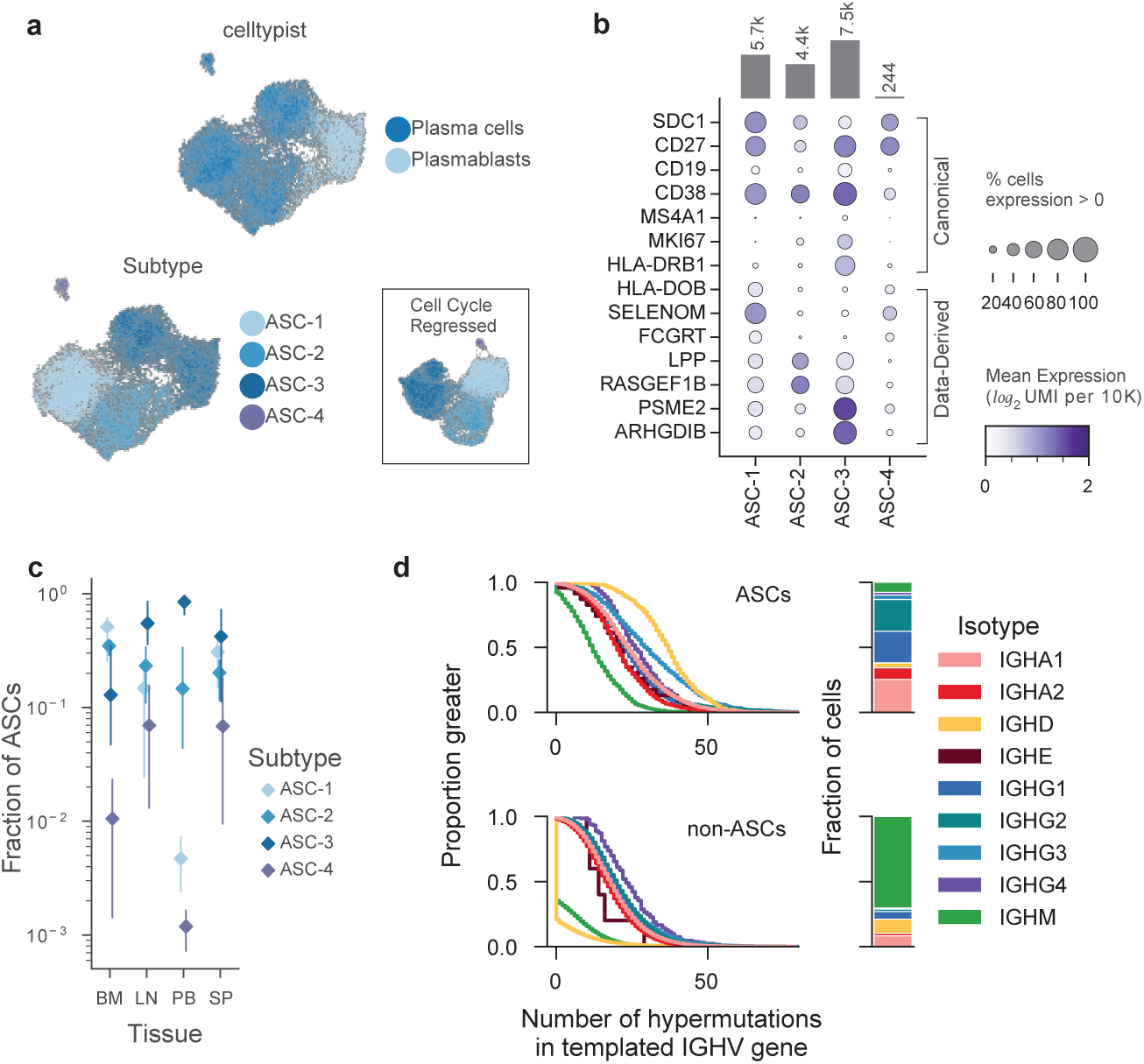
Heterogeneity in ASC transcriptomes and VDJs. (a) Low-dimensional (UMAP) representation of transcriptional heterogeneity in ASCs. Celltypist labels (top), and transcriptome-derived labels (bottom). Inset shows UMAP representation when cell-cycle associated genes are removed. (b) Genes distinguishing between the subtypes of ASCs, where the top half of the plot shows canonical genes often used for flow cytometry and the bottom shows the data-derived, transcriptionally detected genes which distinguish the subtypes. (c) The relative fractional abundance of ASC subtypes in different tissues. Points represent means across all donors and error bars represent the range observed among the different donors. (d) The distributions of hypermutation levels for ASCs and non-ASC B cells of different isotypes (left) and overall isotype usage for ASC and non-ASC B cells.

**Figure ED5.**
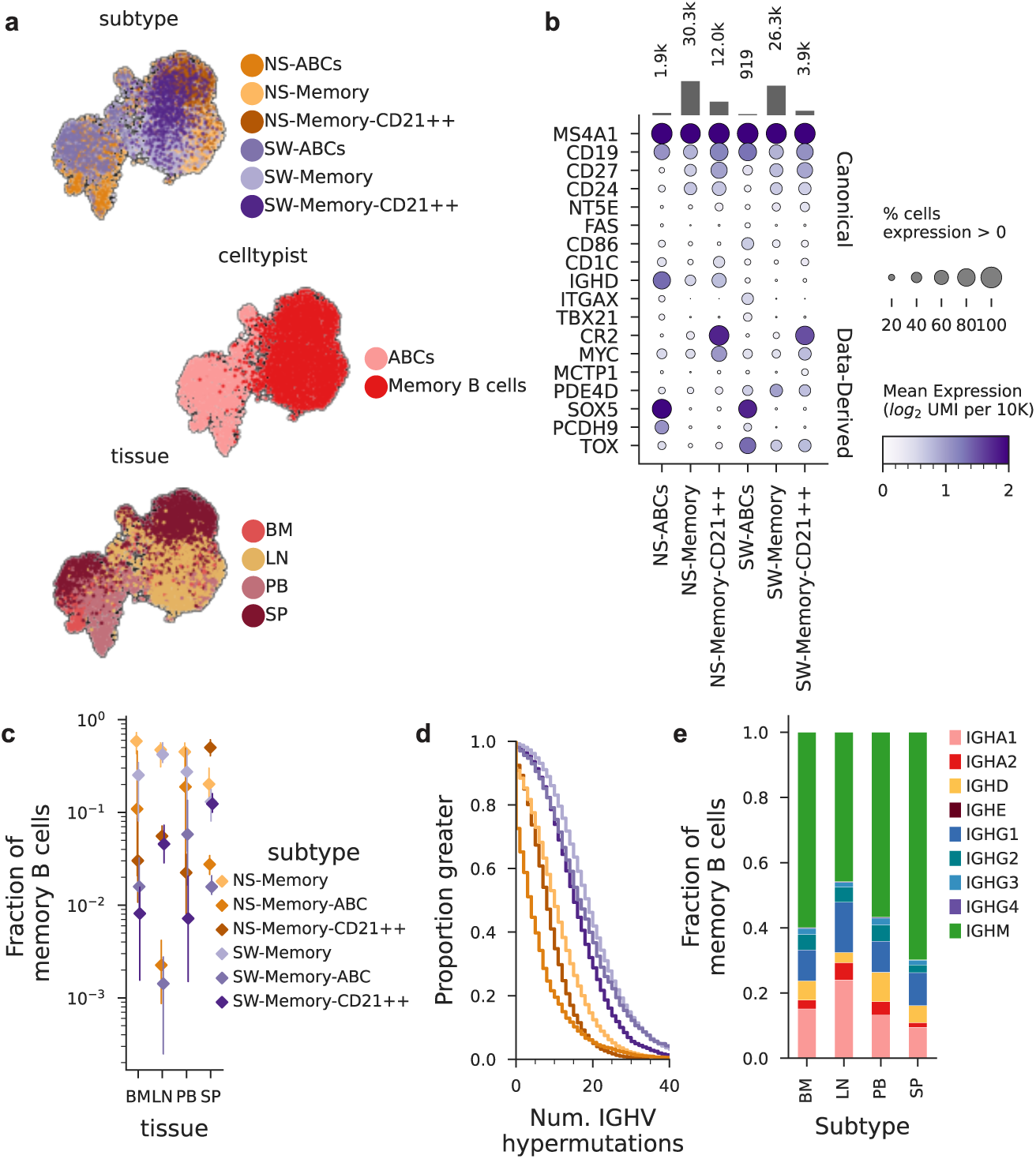
Heterogeneity in memory B cells. (a) Low-dimensional (UMAP) representation of transcriptional heterogeneity in memory B cells. Celltypist labels (top), (middle) transcriptome-derived labels, and tissue (bottom). For these UMAPs, cell types that account for more than 1% of the population were sub-sampled uniformly before calculating the nearest-neighbor graph. (b) Genes distinguishing between the subtypes of memory B cells, where the top half of the plot shows canonical genes often used for flow cytometry and the bottom shows the data-derived, transcriptionally detected genes which distinguish the subtypes. (c) The relative fractional abundance of memory B subtypes in tissues averaged across all donors. Points represent means across all donors and error bars represent the range of the data (d) The distribution of hypermutation levels. (e) Barplot of constant region usage in tissues

**Figure ED6.**
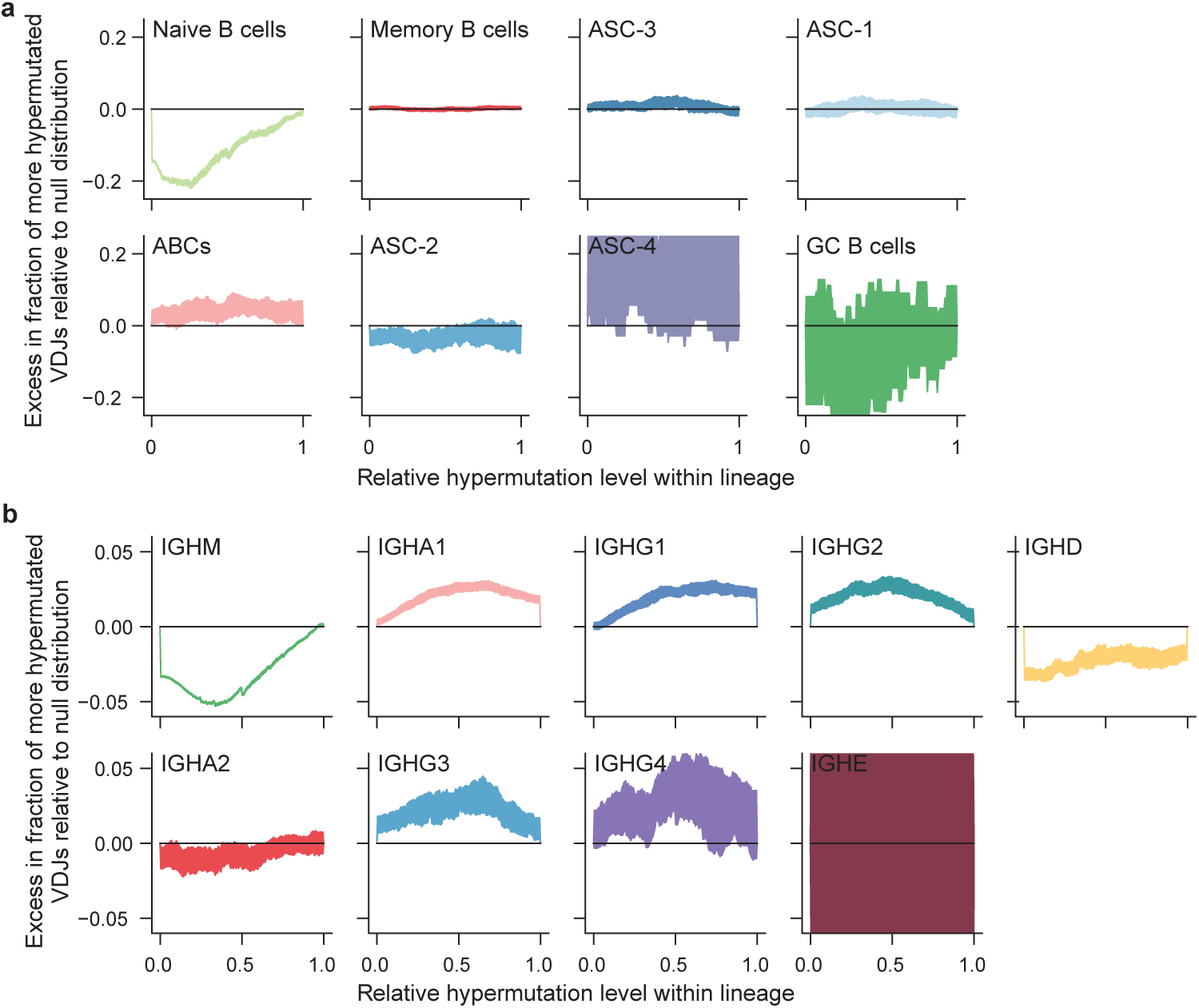
Distributions of cell types and isotypes among cells belonging to the same lineage. (a)The within-lineage distribution of cells of a particular cell type. The panels corresponding to Naive B cells, Memory B cells, and ASC1-3 and duplicated from **Fig. 4b**, and are shown here separately for visual clarity. (b) The distribution of hypermutation levels for all cells in lineages with any cells using a particular constant region. As in **Fig. 4b**, the scaled hypermutation level represents the ratio of the excess in the number of hypermutations relative to the least hypermutated sequence in the lineage, and the excess number of hypermutations of the most hypermutated lineage in the sequence. Null distributions were obtained by permuting IGHC gene labels within lineages containing that isotype. The confidence intervals represent the bounds on the cumulative density function obtained in a 100 independent permutations.

**Figure ED7.**
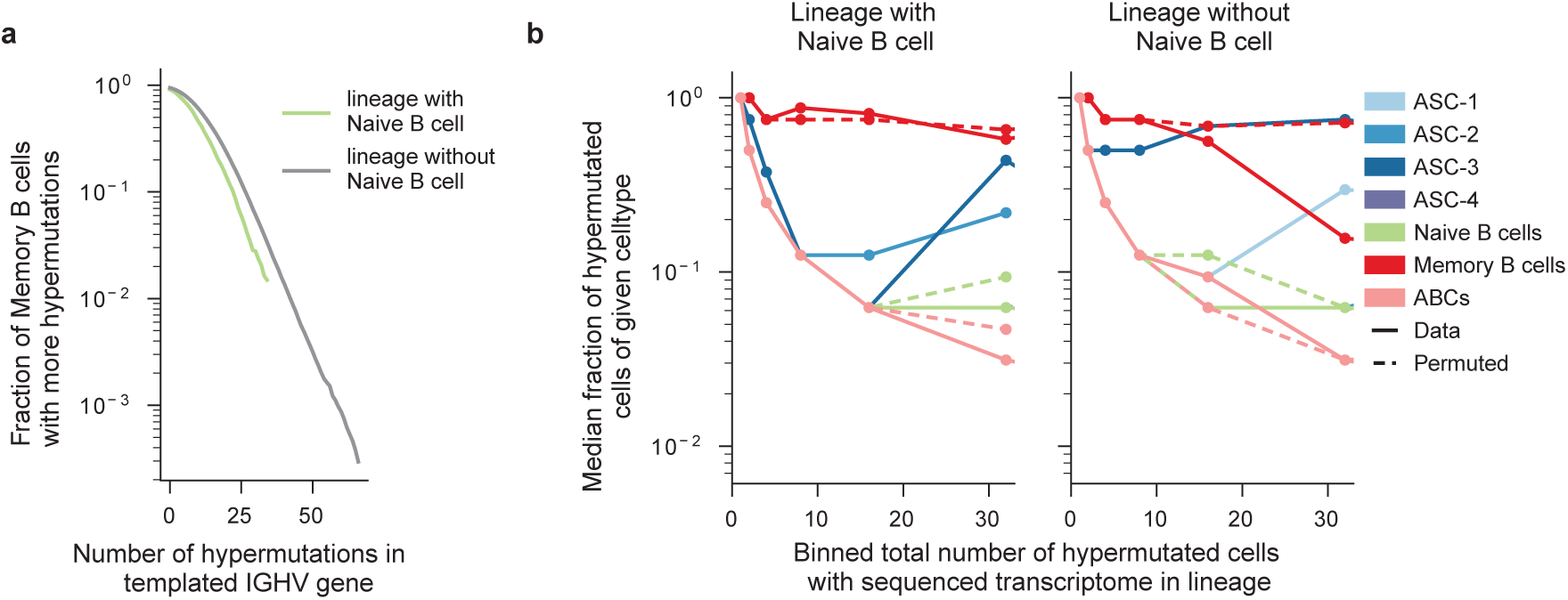
Correlations in cell types of cells belonging to the same lineage. (a) Distribution of the of IGHV hypermuations in all memory B cells (grey) and memory B cells found in the same lineage as an unhypermutated Naive cell (green). (b) Average fraction of cells of given type in lineages in which an unhypermutated Naive cell was sampled (left), and in lineages in which none of the sampled cells were an unhypermutated Naive cell (right). Full lines represent data, and dashed lines represent distributions obtained after permuting cell state labels among all cells with sequenced transcriptomes.

**Figure ED8.**
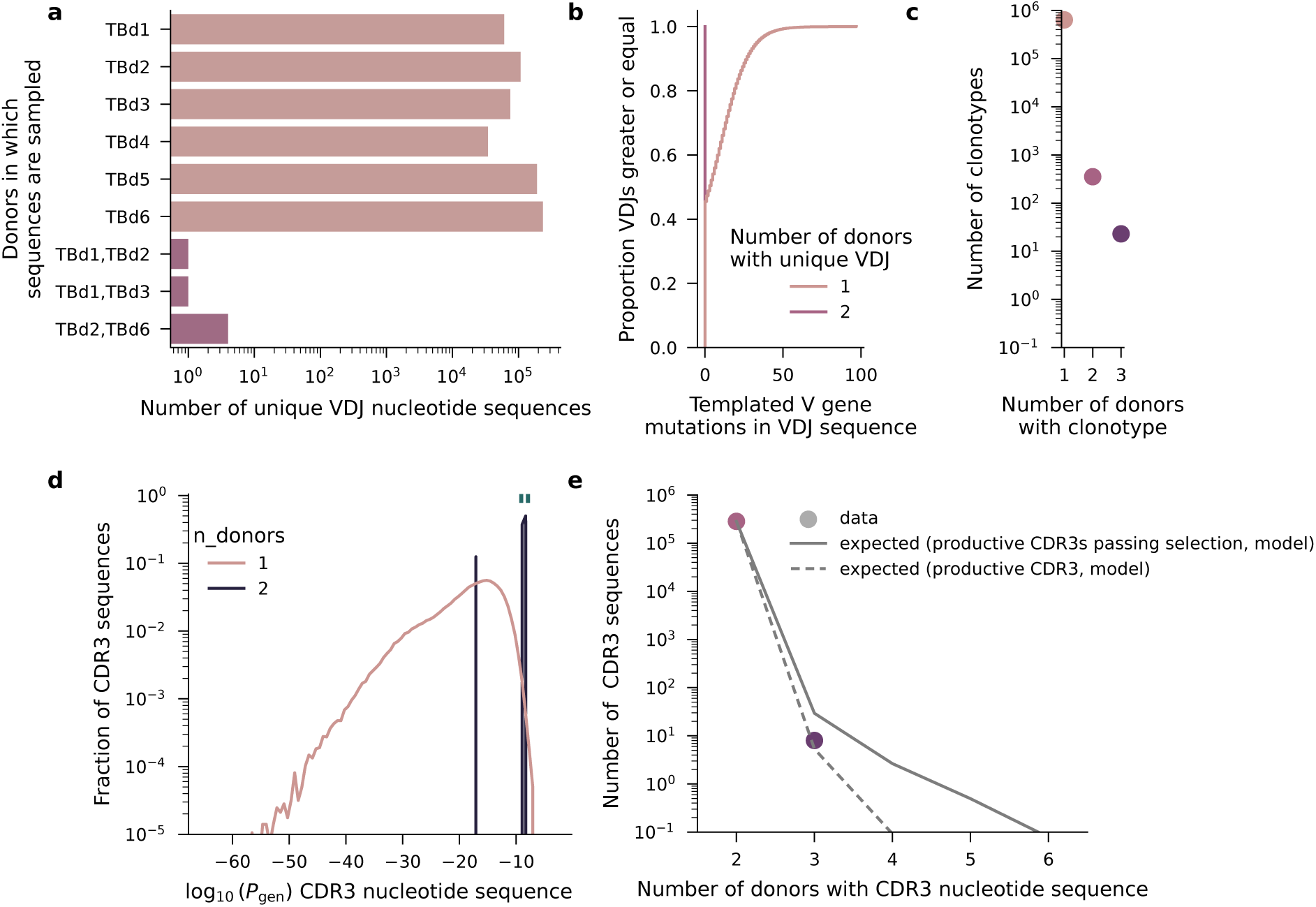
Characteristics of VDJ sequences found in multiple donors. (a) Number of unique nucleotide VDJ sequences as a function of the donors they were found in. (b) Distribution of the number of hypermutations in the templated portion of the IGHV gene as a function of the number of donors a VDJ sequence was found in. (c) Number of shared “clonotypes” in our data, where a clonotype is taken to be a sequence with an identical germline V gene, J gene, and CDR3 amino acid sequence. (d) Recombination probabilities for the CDR3 nucleotide sequence as a function of the number of donors the CDR3 was found in. Green dashes denote the CDR3’s associated with the VDJ sequences shared between multiple donors. (e) Number of shared CDR3s in our data compared to the number expected under given the overall distribution of recombination probabilities shown in (d). The two models are described in detail in the **Supplementary Information Section 4**.

**Figure ED9.**
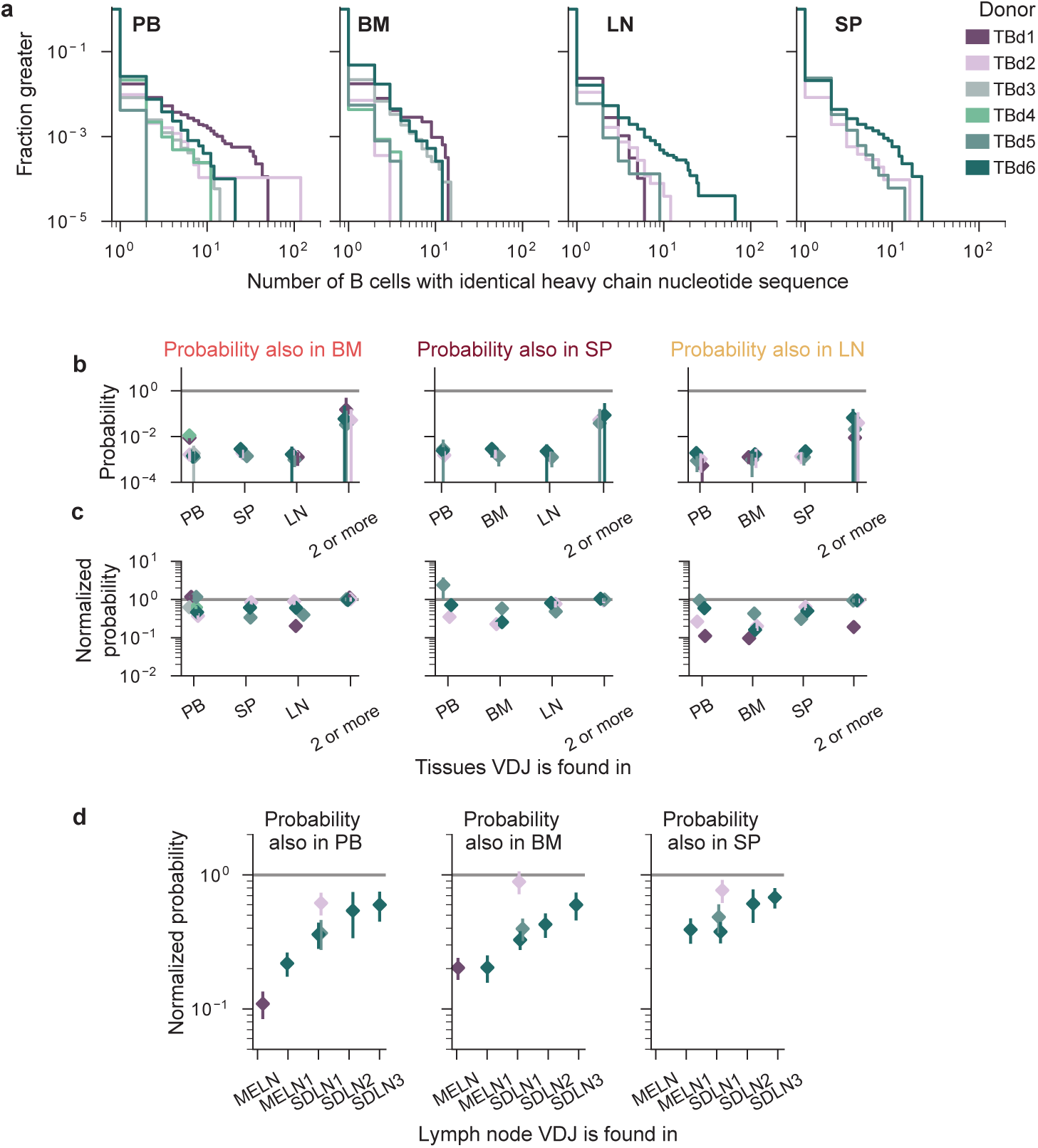
Clonal expansion and hypermutation distributions across tissues and localization statistics of shared B cell clones. (a) To avoid conflation of VDJ dispersion in the ambient and true B cell expansion, the distributions are derived from droplets containing a single VDJ not found in ambient droplets and a high-quality B single B cell transcriptome. (b) Probability that a B cell clone is found in the bone marrow, spleen and lymph nodes, given its VDJ has been sampled in a different tissue, analogously to in **Fig. 6a**. Colors as in panel (a). (c) The normalized probability of appearance in the same tissues, analogously to **Fig. 6b**. (d) Normalized probability that a B cell clone is found in the peripheral blood, bone marrow, or spleen conditional on it having been sampled in a specific lymph node. MELN refers to the mixture of three different mesenteric lymph nodes from TBd1; MELN1 refers to the single mesenteric lymph node from TBd6, and SDLN1-3 refer to the three individual supradiaphragmatic lymph nodes from TBd6.

**Figure ED10.**
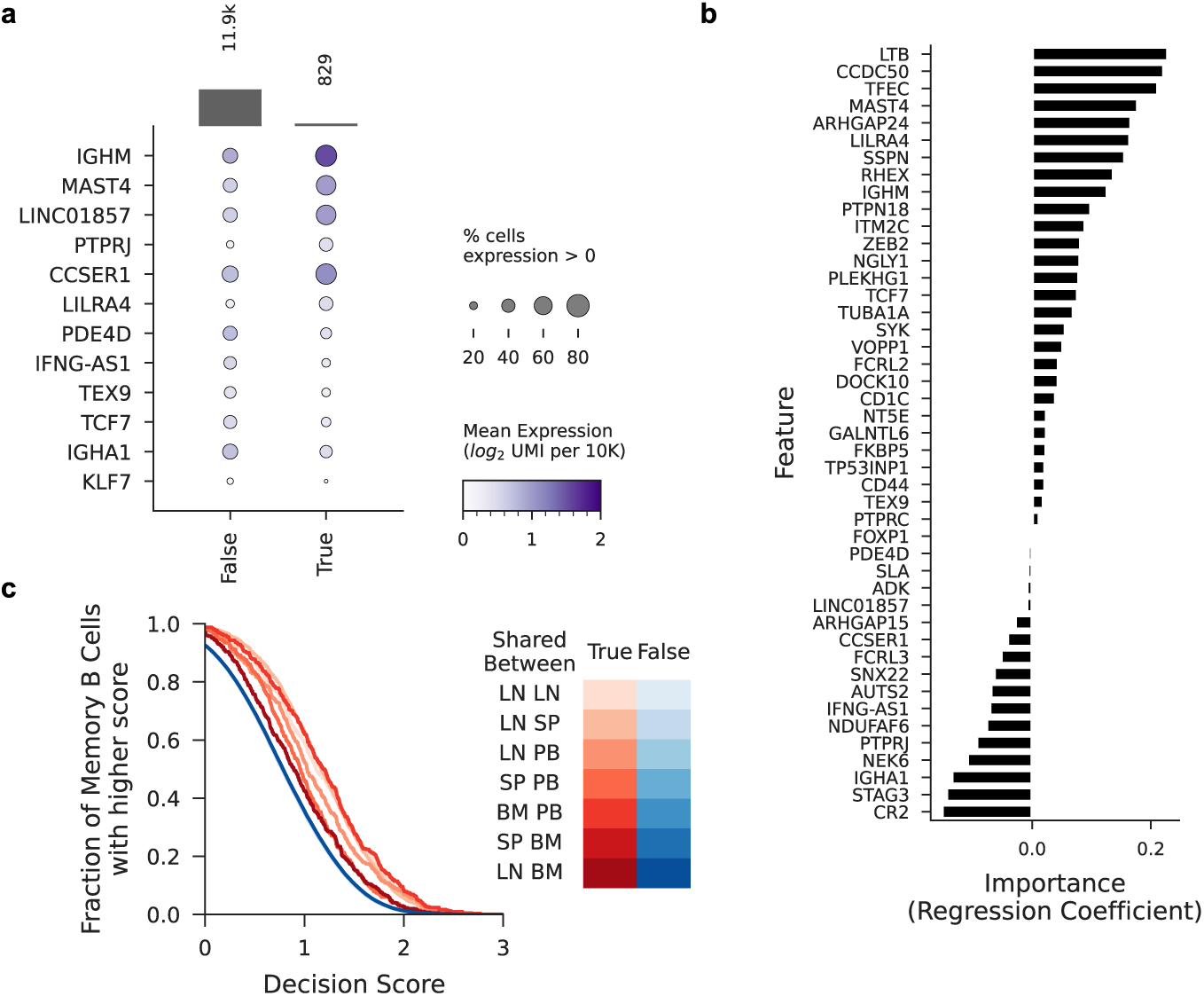
Gene expression signatures of shared memory B cells. (a) Dotplot showing the gene expression differences conditional on whether a memory B cell is shared between Lymph Nodes and Spleen in TBd6 with bars showing total numbers of cells in each group (b) Feature importances for a logistic regression classifier constructed to predict whether or not a memory B cell is shared between Lymph Nodes and Spleen for all donors; features were selected from differential expression analysis on TBd6 shown in (a) (c) Decision scores of the logistic regression classifier for all memory B cells from all donors colored by whether or not a memory B cell is detected as shared between given tissue pair. For details see **Supplementary Information Section 5.3**.

## Supplementary Information

### 1 Experimental Methods

#### 1.1 Tissue processing

##### 1.1.1 Tissue Procurement

Donated organs and tissues were procured at various hospital locations in the Northern California region through collaboration with, Donor Network West (DNW, San Ramon, CA, USA). DNW is a not-for-profit, federally mandated organ procurement organization (OPO) for Northern California. Recovery of non-transplantable organ and tissue was considered for research studies only after obtaining records of first-person authorization (i.e., donor’s consent during their DMV registrations) and/or consent from the family members of the donor. Tissues were processed consistently across all donors. Each tissue was collected, and transported on ice, as quickly as possible to preserve cell viability. A private courier service was used to keep the time between organ procurement and initial tissue preparation to less than one hour. Single cell suspensions from each organ were prepared as described in **Section 1.1.3**. The research protocol was approved by the DNW’s internal ethics committee (Research project STAN-19-104) and the medical advisory board, as well as by the Institutional Review Board at Stanford University which determined that this project does not meet the definition of human subject research as defined in federal regulations 45 CFR 46.102 or 21 CFR 50.3.

##### 1.1.2 Background of Donors

The brief medical history for all donors is outlined in **Table S1**. To be included in our study, donors had to have no history of immune disease or cancer, and no evidence of active infection or pregnancy. Per DNW Management Guidelines, all donors received a number of drugs while on a respirator prior to cross clamp for organ removal. During this time, all donors received antibiotics. Zosyn was the most common, but Diflucan and Vancomycin were given to some donors. Based on culture results, drug allergies or medication availability, other antibiotics were administered to some donors. Donors also received anti-coagulants, anti-inflammatories, diuretics, and drugs to maintain blood pressure. Most commonly these were heparin, Solu-Medrol, Levophed, and Lasix.

##### 1.1.3 Tissue Processing

All 6 donors were processed with standard protocols previously described in Ref. [28]. We reiterate them below for completeness.

###### Blood

We mixed the full amount of blood from the organ donors (between 5 and 40mL) with an equal volume of PBS plus 2% BSA. 15 mL of Density Gradient (Ficoll Histopaque-1119) were added to empty 50-mL Falcon tubes. Up 25 mL of blood/buffer mixture were added to each Ficoll-filled Falcon tube, tilting the tube and pipetting on its side to prevent the blood from mixing with the Ficoll. Tubes were centrifuged at 400g for 30 minutes at room temperature, with the centrifuge brakes off. After centrifugation, the tubes were inspected to check that all layers were well separated. Starting from the bottom, the following layers were identified: erythrocytes, Ficoll solution, buffy coat with cells (white color), and plasma. The buffy coats were gently removed from each tube and transferred into a new 50mL Falcon tube. 30mL of cold (4°C) PBS + 2% FBS were added to each buffy coat. After pelleting the cells in the buffy coat, we washed them twice in cold PBS + 2% FBS, then performed ACK Erythrocyte Cell (ThermoFisher) lysis for 5 minutes. After the ACK lysis step, cells were washed with 10mL of ice-cold PBS and spun down at 4°C, 500g for 10 minutes. Cells were then resuspended in buffer and enumerated using a hemocytometer.

###### Bone Marrow

The vertebral bodies (VB) were wrapped in a saline-soaked cloth and shipped to Stanford University on ice. Upon arrival, the VBs were cleaned of connective tissue and fat using sterilized wood chisels. VBs were then cut into ∼ 2cm^3^ pieces using bone cutting forceps and rongeurs. The bone marrow pieces were transferred into a plastic container, to which 100µL of RPMI + 10% FBS was added. The container was mechanically tumbled for 30 minutes at room temperature. At this point the buffer was turbid and cellular. This was then passed through a 100 µmstrainer into 50mL falcon tubes which caught bone chips and other smaller debris. Multiple strainers were often used due to clogging. After straining, the cells were centrifuged and pelleted at 330g for 5 minutes at 4°C. Cells were then washed twice with PBS + 2% FBS andresuspended in PBS + 2% FBS. BMMNCs were then isolated using Ficoll by layering 35mL of cell suspension on 15mL of Ficoll density gradient medium. Cells were centifuged at 445g for 35 minutes at 20°C in a swinging bucket rotor without braking. Mononuclear cells were transferred into a new 50mL Falcon tube. 30mL of cold (4°C) PBS + 2% FBS were added to each buffy coat. After pelleting the cells in the buffy coat, we washed them twice in cold PBS + 2% FBS, then performed ACK Erythrocyte Cell lysis for 5 minutes. Cells were then centrifuged for 5 minutes at 330g at 4°C without brakes. Cells were then resuspended in PBS + 2% FBS and enumerated using a hemocytometer.

###### Spleen

The spleen tissue was placed in a petri dish and minced with sterile surgical scissors and scalpels. The minced tissue was transferred to a 50mL conical tube with 5mL of digestion media, which was freshly prepared (0.8 mg/mL Collagenase IV (Worthington) and 0.05 mg/mL DNase I (Roche) in RPMI with 10% FBS). The tissue was further minced with scissors inside the 50mL tube. The petri dish containing the tissue was washed with 5mL of digestion media, and everything was transferred to the 50mL tube. The tissue was digested in a shaker at 37°C, 200 rpm for 30 minutes. The sample was vortexed every ten minutes to re-disperse the tissue. After digestion, we pipetted the mixture vigorously to evaluate digestion and to further mechanically digest the tissue as much as possible. The tissue solution was diluted in RoboSep buffer (Stemcell Technologies) and then passed through a 100µm cell strainer into a 50mL tube. A plunger from a 10mL syringe was used to mash the remaining tissue in the strainer. The cell suspension was spun at 4°C, 450g for 10 minutes and washed twice in RoboSep buffer. Finally, we performed ACK Erythrocyte Cell lysis for 5 minutes, after which cells were washed with 10mL of ice-cold PBS and spun down at 4°C, 500g for 10 minutes, then enumerated with a hemocytometer.

###### Lymph Nodes

Individual lymph nodes were collected in a p100 petri dish and the surrounding edges were cleaned to extract lymph nodes from surrounding fat, using sterile surgical scalpels and scissors. Once the vast majority of fat was removed, the lymph nodes were placed in a 5mL polypropylene tube and minced with sharp scissors. Digestion media was prepared with 0.8 mg/mL Collagenase IV (Worthington) and 0.05 mg/mL DNase I (Roche) in RPMI plus 10% FBS. Up to 5mL of digestion media was added to the 5mL polypropylene tube, which was then placed in a shaker to digest at 37°C, 200 rpm for 20 minutes. At the end of the 20 minutes, the cell suspension was pipetted vigorously up and down 10 times to evaluate digestion and to further mechanically digest the tissue as much as possible. The tissue solution was diluted in RoboSep buffer and then passed through a 100µm cell strainer into a 15mL tube. A plunger from a 10mL syringe was used to mash the remaining tissue in the strainer. Cells we spun down and resuspended in RoboSep buffer. After an optional ACK lysis step (based on whether the pellet looked red), cells were spun down at 4°C, 450g for 10 minutes, resuspended in 5mL of buffer and enumerated with a hemocytometer.

###### Cell freezing and thawing

For freezing, cells were spun down and resuspended with around 200*µ*L of buffer. Ice-cold Cryostor CS10 buffer was added to the cells to attain a density between 10 and 40 million cells per mL. The cell suspensions were transferred to cryovials, and frozen using a standard slow rate-controlled cooling protocol (approximately -1°C/minute) in CoolCells. in a -80°C freezer. After a few hours cells were moved to long-term storage in liquid nitrogen vapor. For thawing, we followed the 10X Genomics thawing protocols, which appear to differ from typical protocols only in that they add RPMI + 10% FBS (or similar) at a slower rate. Initially, we experienced catastrophic amounts of cell aggregation, in particular in bone marrow cells, which was mitigated by using 50U/mL of Benzonase (Sigma) in the thawing media.

###### B cell purification and loading on Chromium or Chromium X

Cell suspensions were counted and resuspended at target densities of 5 × 10^7^ cells per mL. B cell purification was performed according to manufacturer instructions with the Stem Cell EasySep Human Pan-B Cell Enrichment kit (Stemcell Technologies). After enrichment, cells were resuspended in RoboSep Buffer and counted. We loaded cell suspensions on the Chromium or Chromium X (10X Genomics) instruments according to manufacturers instructions, except when performing VDJ only profiling. We targeted 20,000 recovered cells per lane when measuring gene expression and VDJ profiling and 80,000 recovered cells per lane when performing VDJ only profiling.

#### 1.2 Transcriptome and VDJ sequencing

Gene expression libraries were generated following the 10X manual. VDJ amplicons were amplified and enriched by a nested PCR using the same primers as quoted by the 10X manual, but in custom PCR mixes, containing KAPA 2X HotStart (Roche), the 10X forward VDJ amplification primer at a final concentration of 1µM, and the pool of 10X isotype-specific reverse primers at a final concentration 0.5µM each and in a total volume of 50µL. Both the inner and outer target enrichment PCRs were performed for 8 cycles, annealing at 67°C. We then proceeded to make libraries from these amplicons according to the 10X manual. Individual libraries were dual-indexed and sequenced on Illumina NextSeq 2000, or Illumina NovaSeq S2 or S4 flowcells. We targeted a sequencing depth of 5000 reads per cell for VDJ amplicons and 40 000 reads per cell for the gene expression libraries.

We noted that antibody secreting cells express on average multiple orders of magnitude more antibody transcripts than naive or memory B cells, which is unsurprising given the majority of detectable RNA molecules in antibody secreting cells are derived from the IGH and IGK/L loci[43]. Specifically, whereas the reconstructed IGH amplicons for most B cells in our study were supported by fewer than 20 IGH UMIs (**Fig. S1**, bottom panel), 1-10% of cells in each sample were supported by thousands or even tens of thousands of UMIs. As a result, the majority of captured transcripts, and thus the majority of captured reads, is associated with these cells. This results in the preferential detection of antibodies of ASC origin in our sequencing libraries, and the corresponding loss of VDJ transcripts associated with memory or naive B cells, which have vastly lower expression.

To understand whether, in principle, sequencing more deeply would allow us to detect additional antibody transcripts from naive and memory B cells we examined what fraction of UMIs were available to the typical cell in our samples (**Fig. S1**, top panel). We found that in all samples the fraction of total molecules sequenced from ASCs was over 90%. In certain samples, a small number of ASCs accounted for almost 999 in 1000 reads, with 99 in 100 being the more typical rate. Thus, even when targeting 5000 reads per cell, this effectively allocated no more than 5 to 50 reads to the median memory or naive B cell, causing them to frequently drop out from our libraries. As a result, we concluded that full recovery of our naive and memory B cell transcripts would require us to target about 500,000 reads per cell in the VDJ amplicon libraries, a depth currently practically infeasible given the scale of our study and the cost of sequencing.

**Figure S1:**
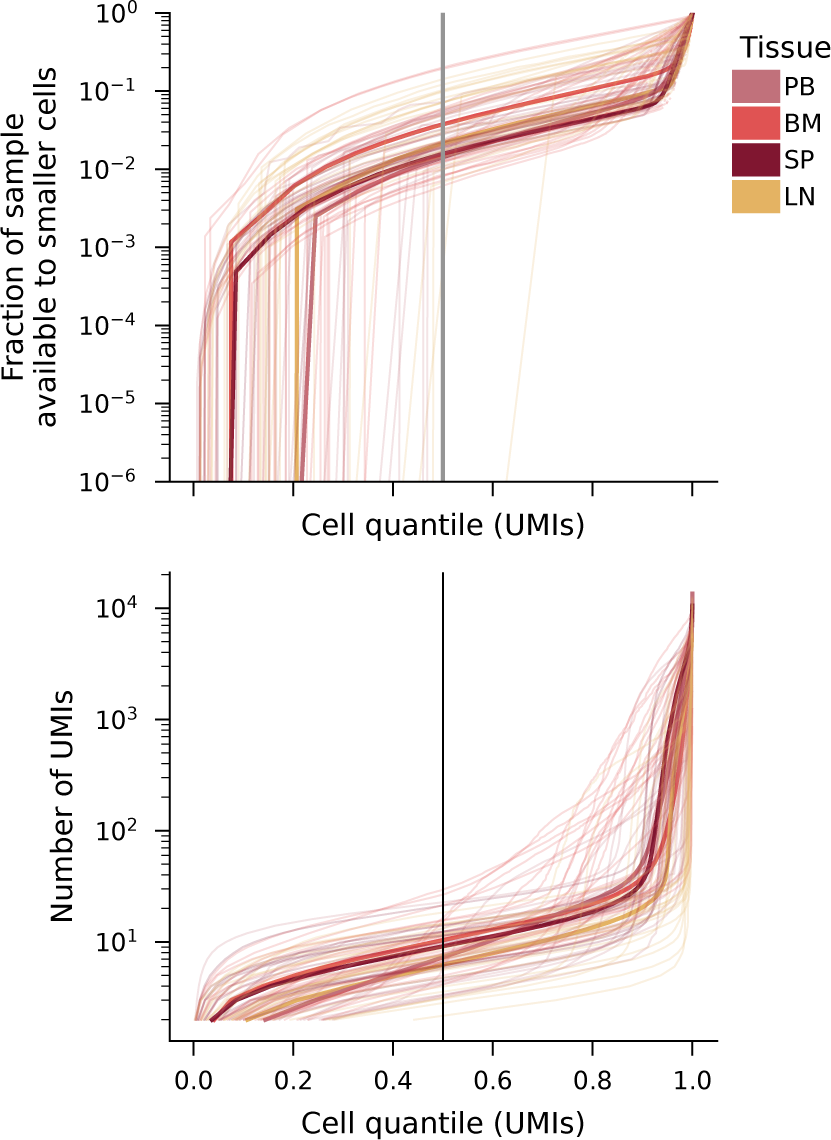
Sample and read availability and depletion by antibody-secreting cells. Number of UMIs (bottom) and cumulative fraction of sample available (top) as a function of cell IGH expression quantile. Thin lines correspond to individual libraries, thick lines represent the aggregate distributions for all samples of a certain tissue. Individual samples and aggregate distributions are colored by tissue of origin.

We note that this problem is fundamental to ASC presence in a mixed library, and arises in all circumstances in which these cells are not separated from memory and naive B cells prior to lysis. As such, it persists when sequencing antibody transcripts in bulk. Thus, we recommend that researchers physically separate cells with ASC phenotypes from memory and naive B cells, prior to lysis, to ensure the balanced recovery of transcripts from each of these subsets.

#### 1.3 Complete enumeration of tissue samples used in the study

**Table S2:**
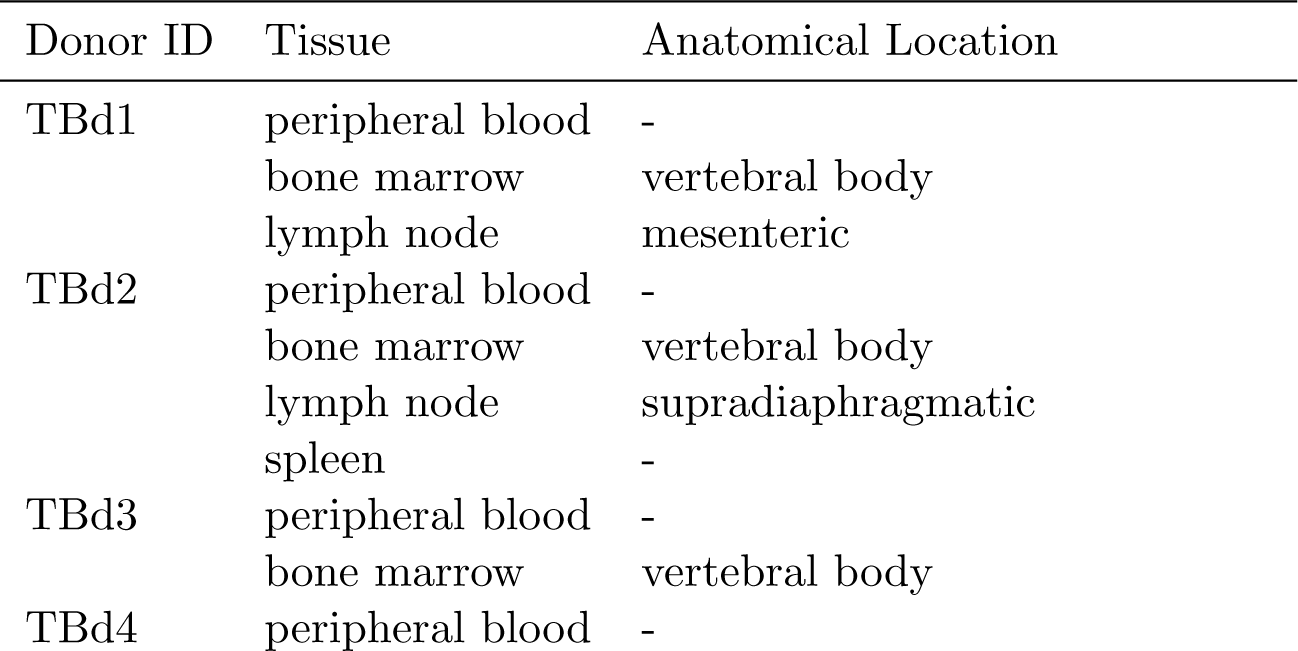
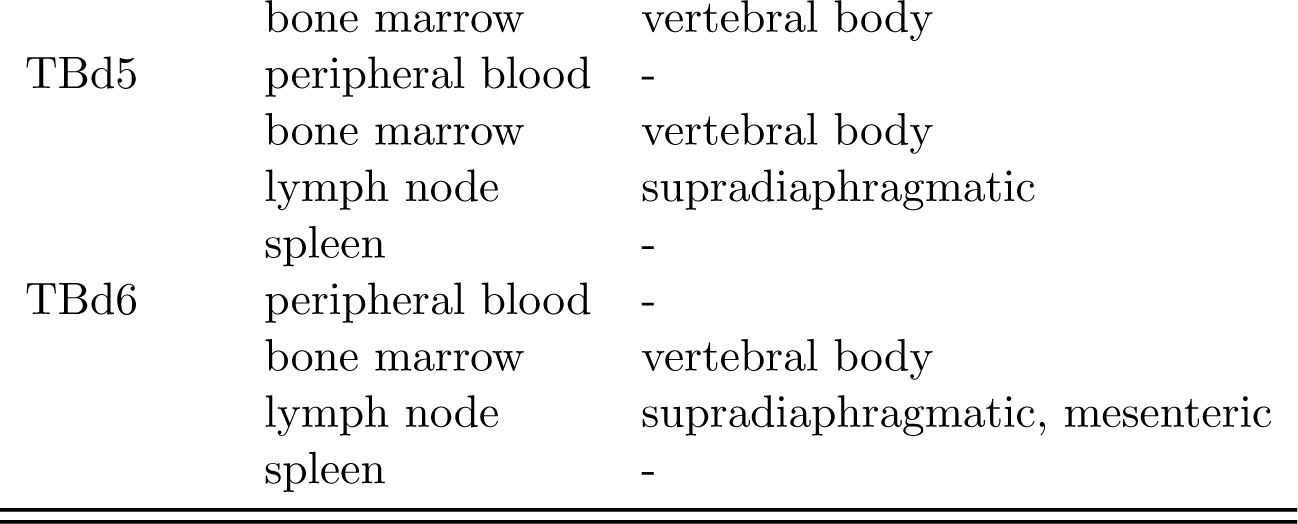
List of all tissues used in this study and their anatomical locations.

#### 1.4 Complete enumeration of cell suspensions, 10X lanes, and libraries sequenced

**Table S3:**
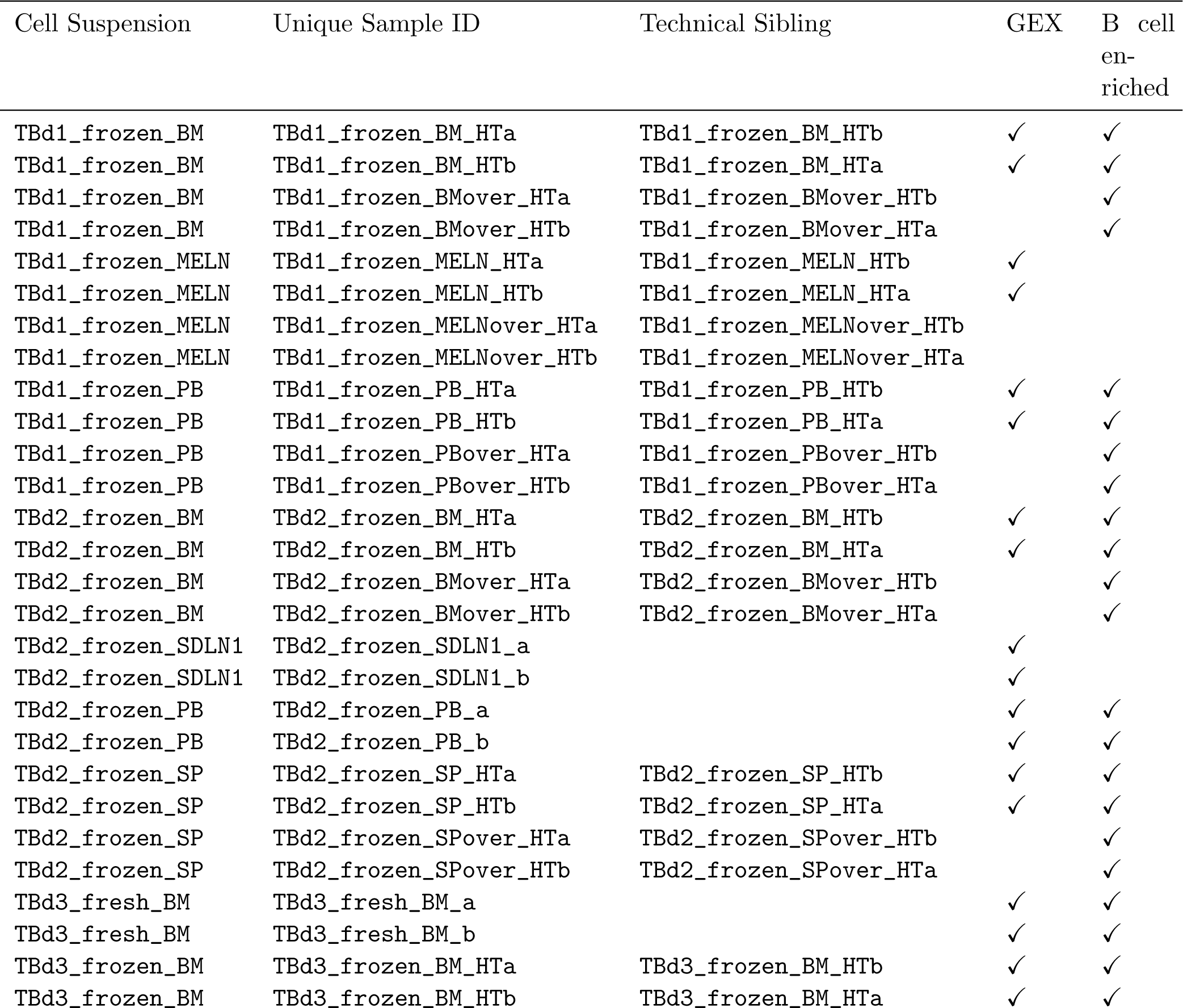
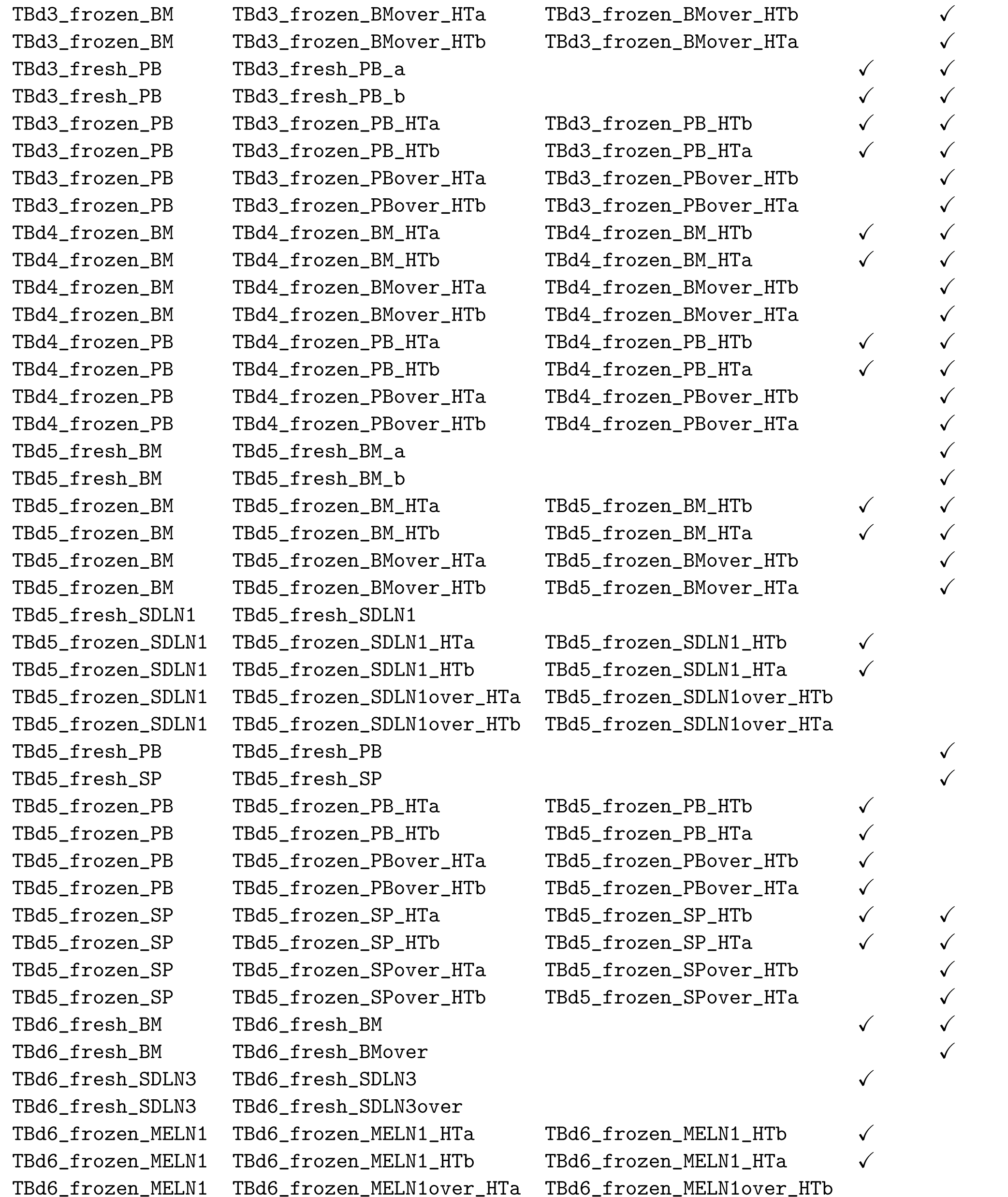
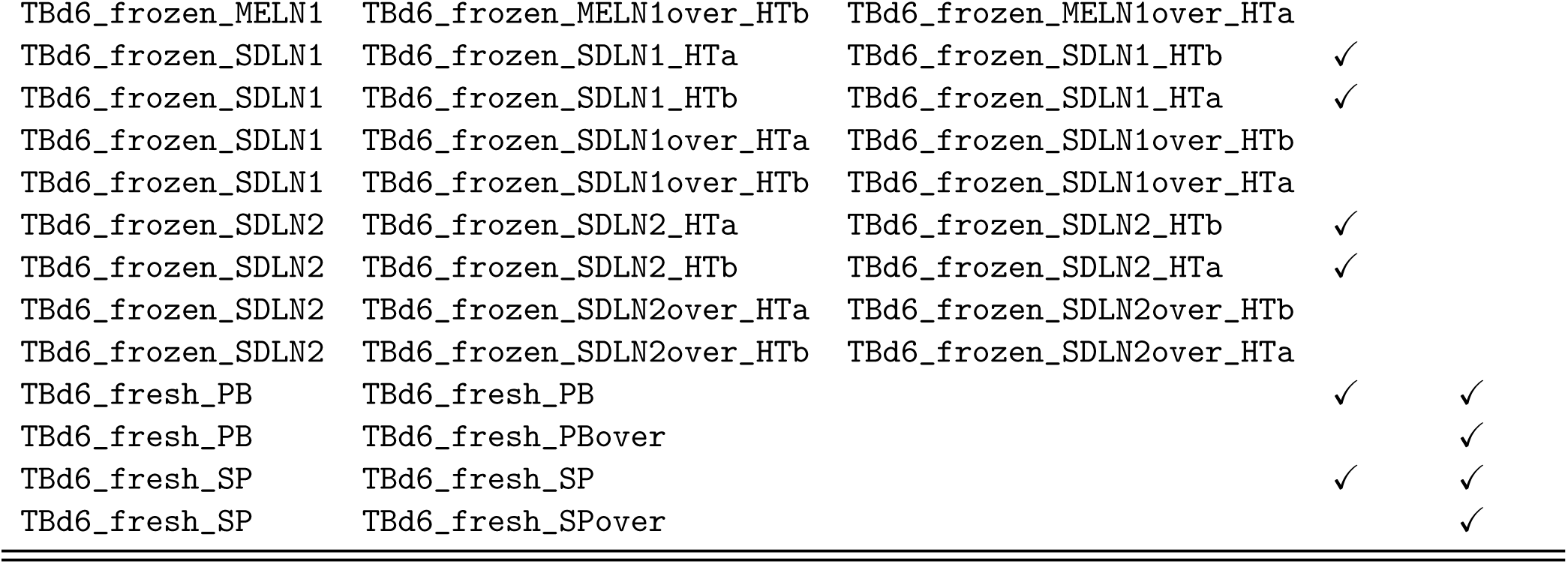
Enumeration of all sequencing libraries analyzed in this study. VDJ libraries were prepared and analyzed for all samples listed. Samples have HT in their name if they were generated on the 10X Chromium X Controller; all other samples were generated on the 10X Chromium Controller. Unless designated as technical siblings, samples of the same tissue represent separate samples of the live cell suspensions noted in the column. Technical siblings represent the two equal volume parts of the same emulsion, generated on the 10X Chromium X Controller using the 5*^′^* high throughput kit. These were typically processed, indexed, and sequenced separately. We did not split the emulsion in samples generated on the standard throughput kit on the Chromium Controller. “B cell enriched” denotes that negative selection for B cells was performed on the sample.

### 2 Gene expression analysis

#### 2.1 Gene expression data preprocessing

We used `cellranger v7.0.1` to align reads and count UMIs from the raw FASTQ gene expression data. Reads were aligned to the GRCh38 human reference genome. The unfiltered outputs of `cellranger` were used as inputs to `cellbender`[44], which develops a per-sample model of ambient RNA and produces a count matrix that has removed putative ambient RNA. While we used these count matrices for orientation in exploratory analysis, all cell type annotation and gene expression analyses were ultimately based on the raw UMI counts. Cell barcodes with fewer than 1000 UMIs or with fewer than 500 genes detected were excluded from downstream analysis. To facilitate doublet-detection, understand batch-effects, and gain additional statistical power for cell type identification, we added the entire dataset from Ref. [7] to our pipeline before training scVI models. The dataset was downloaded from the Ref. [45]. Pan Immune Dataset cells were not included in any of the analyses, statistics and visualizations presented in the manuscript.

#### 2.2 Doublet detection and automatic annotation

We performed doublet detection on a per-sample basis using `scrublet` [46]. We noticed the algorithm likely had a high proportion of negatives, given the inferred doublet rates were often far “faction-voting” methods and one gene-based heuristic to flag cells as doublet-associated, regardless of their doublet score.

First, we flagged cells if their gene expression signature is similar to that of other cells with high doublet scores. We labeled these cells “doublet-associated”. Doublet-association is defined as at least 1 in 10 of the cells in a given Leiden cluster are called doublets by the Scrublet algorithm, all cells in that cluster are flagged as doublet-associated.

Second, we flagged cells based on `celltypist` labels for their cluster. Using the “Immune All High” model, we predicted broad cell type labels for each cell (e.g. “B cell”, “Macrophages”). We then clustered all cells using (`sc.tl.leiden(resolution = 3)`). Clusters with mixed membership (i.e. with a minority member fraction *>* 0.1) were also flagged as doublet-associated by cell voting. While this heuristic generally performed very well, cycling cells such as plasmablasts were often flagged as doublet-associated. We believe this is due to their strong cycling signature causing them to cluster with cycling cells of other immune lineages. Thus, we exempted cycling cells from this flag. Finally, using gene expression signatures of observed non-B cell contaminants, we created contaminant gene scores `sc.tl.score_genes()` using MPO, AZU1, ELANE, and S100A8 as the Myeloid score, and CD3E, CD3D, and CD247 as the T cell score. For subsequent integration with VDJ contig data, we labeled cells as probable high-quality single B cells if:

- `doublet_score` is less than 0.01,
- `t_cell_score` is less than 0,
- `myeloid_score` is less than 0,
- `celltypist` identifies the cell as a B cell
- `celltypist` confidence score is *>* 0.95

#### 2.3 Cell cycle assignment

We used MKI67 as a marker of cell cycle status. However, we noted that in cell types we often did not detect MKI67, even in plasmablasts which are thought to be cycling (**Fig. S2**, upper left). This is likely due to technical limitations in sensitivity via droplet-based RNA sequencing. Thus, we took advantage of the co-variation between cell cycle genes to generate a more sensitive cell-cycle classification. First we calculated the correlation coefficient of all genes with MKI67. The distribution of the correlation values had a long tail of hundreds of genes with high (*>* 0.5) correlations. Manual inspection of the available evidence for the functions of these genes showed they were mostly involved in the G2/S phases cell cycle. We then used the top 30 most correlated genes to as a set create a cell cycle score using `sc.tl.score_genes()`. This score separated cell types known to be cycling from those that were not with dramatically better sensitivity and specificity (**Fig. S2**, upper right). We validated this score on a dataset of cycling B cells from Ref. [34], which we used to derive a threshold for calling cycling B cells (**Fig. S2**, bottom panels).

#### 2.4 VAE model training and batch alignment

We used scVI [47] to build a model of gene expression for all cells in the dataset based on their raw UMI counts. We found scVI performed batch correction more effectively than other methods we explored [48], at the expense of increased computational resources taken to train the models. We used the default `scvi.model.SCVI` parameters except we changed the number of latent dimensions to 30 and the number of layers to 2. We explored the use of many different batch keys, and ultimately used the concatenation of the “donor” identifier and “tissue” label of each sample as a batch key. We also trained scVI models which added nuisance genes such as the top cell cycle genes, IGH genes, known tissue dissociation genes. All of these latent representations of cells were used for exploratory data analyses in Scanpy, to understand how clustering results where affected by these variables. Ultimately, with the exception of the UMAP in (**Fig. ED4a**) where cell cycle is regressed, all neighborhood graphs were calculated using the scVI model trained on every cell in the dataset and the combination of “donor” and “tissue” as a batch key.

**Figure S2:**
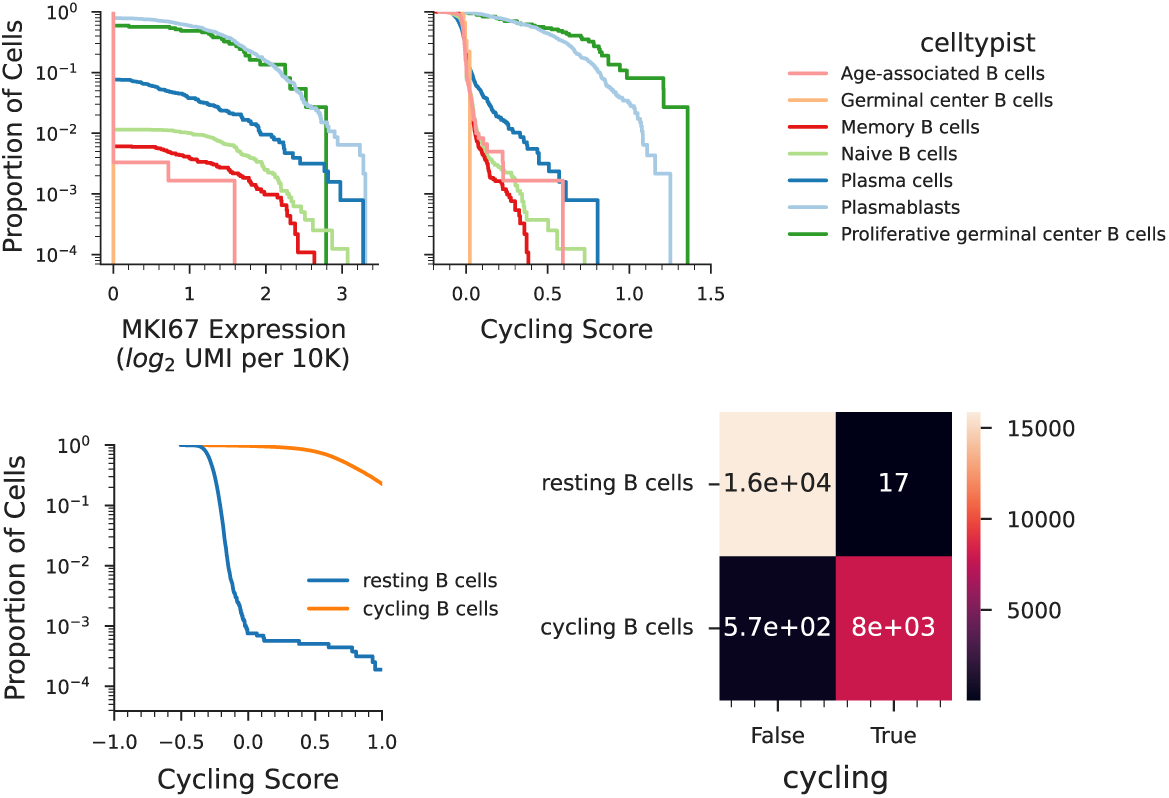
Cell Cycle Annotation. (Top) Distributions of MKI67 expression (left) and Cell Cycling scores (right) colored by celltypist labels. (Bottom) Distributions of Cell Cycling Scores for *in vitro* cycling B cells and resting B cells for [34] (left) and confusion matrix showing the ability of cycling score to distinguish between cycling and non-cycling B cells from at a score cutoff of 0.01

#### 2.5 Biases in cell type counting

Our B cell enrichment was performed via negative selection on all samples except samples derived from lymph nodes (in which B cells account for more than 30% of all cells even in the absence of enrichment) and two TBd5 peripheral blood samples, which we did not enrich because we had a very small amount of starting material for this donor and tissue. While this approach afforded an essentially unbiased look at the relative abundances of different B cells, it resulted in a relatively impure product particularly in bone marrow samples (**Fig. S3**, left panel). We noted that many of the contaminants were progenitor hematopoietic cells (∼ 17%) which may have escaped negative selection because of lower expression of their lineage markers.

When we merged VDJ data with gene expression data on a per cell basis, we noted that many high quality B cell transcriptomes did not have VDJ associated with them. This appears to be due to dropout of VDJ transcripts from low-expressing B cells such as Memory B cells (see also **Fig. S12** and the discussion in **Section 1.2**). Thus, the set of VDJ transcripts that we captured is inherently biased towards antibody-secreting cells (see **Fig. S3**, right panel). As discussed in **Section 1.2**, we anticipate that this bias is present in all unsorted B cell VDJ sequencing. We design all analyses that follow in such a way that they are minimally sensitive to this bias.

**Figure S3:**
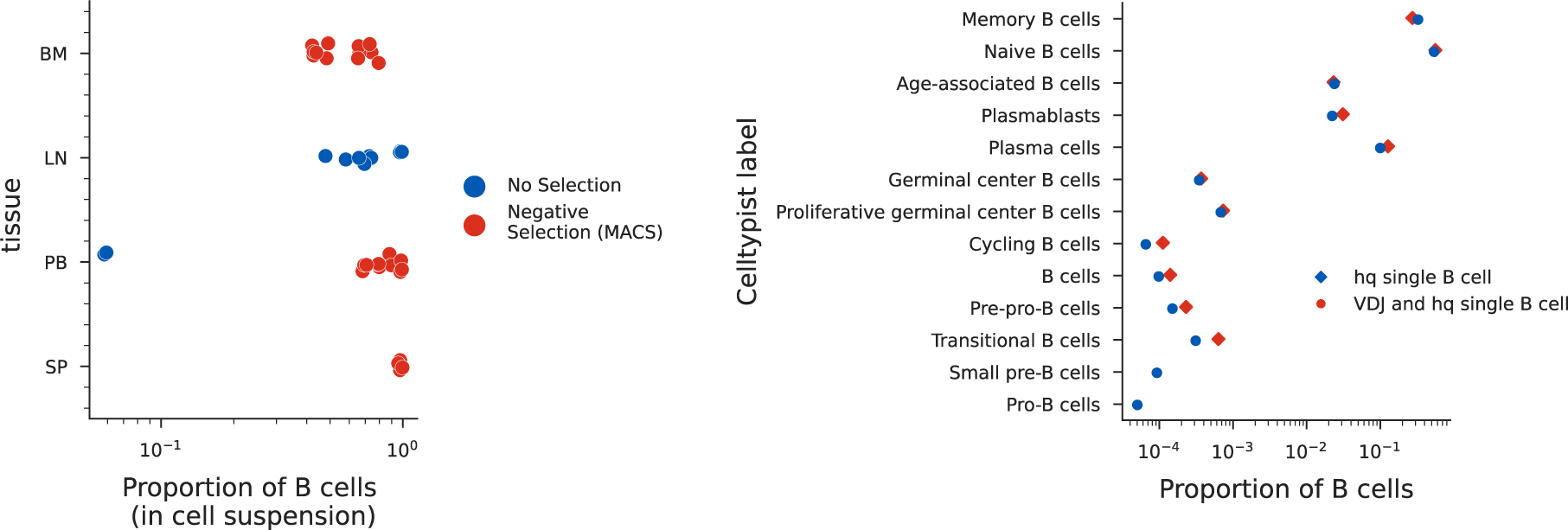
Biases in relative abundances of B cell types during sampling. (Left)Fraction of all cells that are B cells in each cell suspension. Samples are colored by whether or not a B cell purification was performed. (Right) Relative abundances of B cell subsets when either conditioned on having an associated VDJ or not. Cell type labels are from the celltypist Immune All Low model.

Finally, we investigated whether cryopreservation of samples leads to biases in B cell subtype composition, i.e. whether different subsets of B cells survive the freeze-thaw at different rates. To assess this, we prepared replicates of the peripheral blood and bone marrow cell suspensions from donor TBd3, the first immediately after receipt, the second after cryopreservation and thawing. As can be seen in **Fig. S4**, ASC-3s appear most prone to loss during the freeze-thaw procedure with an at least 3-fold survival deficit compared to other B cell subsets, followed by ASC-2s, with no meaningful changes to the abundances of other cell types (for details of the fine-grained annotation of the mentioned cell types, see **Section 2.6**).

**Figure S4:**
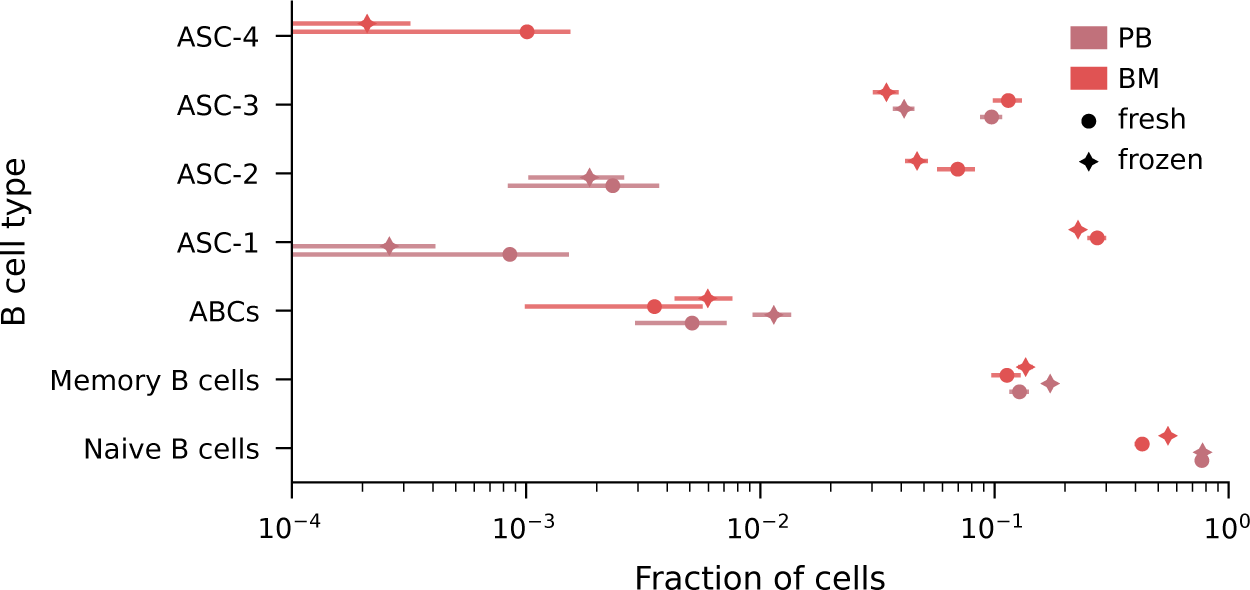
Biases in cell subsets due to the freeze-thaw procedure. Proportion of high-quality singlet non-ambient B cells in each B cell subset of the peripheral blood and bone marrow of TBd3. Error bars denote 95% confidence intervals obtained under the assumption of binomial sampling.

#### 2.6 Fine-grained annotation of B cell subtypes

High-quality single B cells were separated into Memory, Antibody Secreting Cell, and Naive types using the `Immune_All_Low` model and analyzed separately. While we explored a variety ways to analyze these gene expression profiles, ultimately we used the scVI representation to construct nearest-neighbors graphs (`sc.pp.neighbors`), which were clustered using the Leiden algorithm at resolutions between 0.5 and 1 (`sc.pp.leiden`). For ASCs, used an scVI representation with MKI67, RRM2, and TK1 supplied as continuous covariate keys to remove the contribution of the cell cycle. The resulting representation collapsed the distinction in the nearest-neighbors graphs between a particular cluster of Plasmablasts and Plasma cells (ASC-3) (**Fig. ED4a**).

For memory B cells, we noted two major axes of gene expression variation beyond what is described by `celltypist` (**Fig. ED5**). The first was explainable by class-switch status and the second axis was defined by CR2 and CR1 expression (**Fig. S5a**). Thus we used these genes to calculate a Complement Receptor score (`sc.tl.score_genes(adata, gene_list = [‘CR2’, ‘CR1’]`) and defined cells with a positive score as CD21++ (**Fig. S5b**). We incorporated antibody constant region information from the VDJ sequencing data by classifying cells as being switched (SW) if their assembly was not IGHM or IGHD, and non-switched (NS) otherwise. These two simple heuristics captured most of the detectable variability in B cell subsets, and agreed well with Leiden clustering results (**Fig. S5c**).

**Figure S5:**
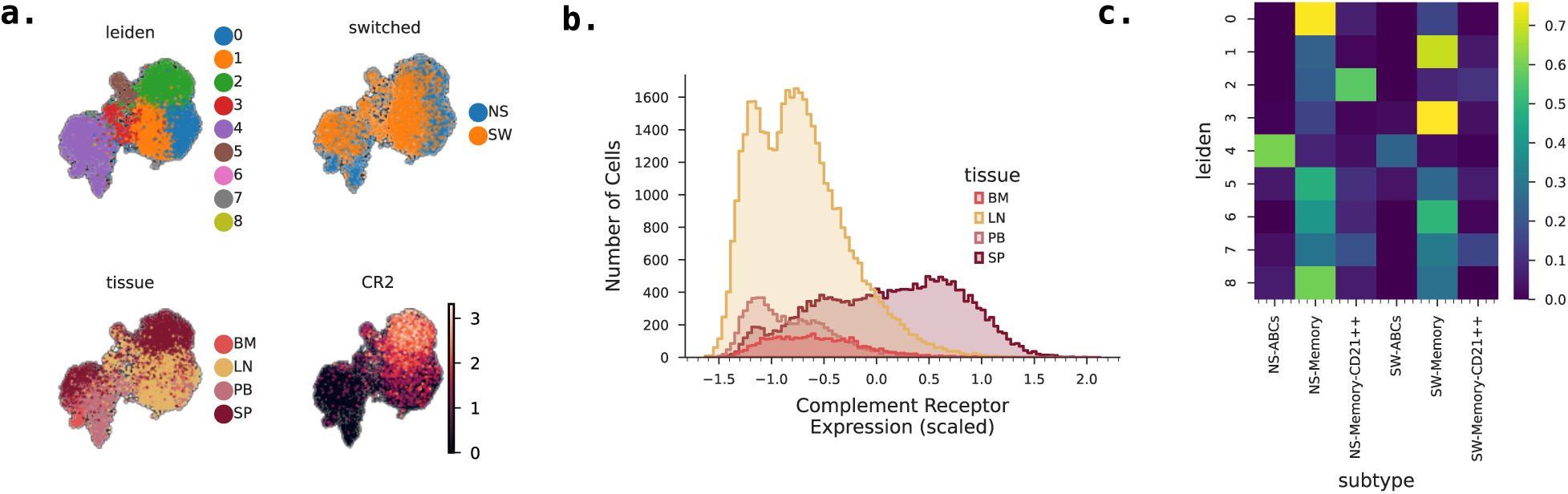
Memory B cell phenotypes. (a) UMAPs showing categorical axes of variation in memory B cell subsets (b) Distributions of complement receptor expression derived scores for memory B cells colored by tissue (c) Confusion matrix showing the agreement between leiden-based data-dependent clustering and heuristic clustering

### 3 VDJ sequence analysis and cell calling

#### 3.1 VDJ sequence preprocessing

We used `cellranger v7.0.1` to filter, trim and assemble VDJ reads into contigs. Our study design included both high-density loading of the VDJ -only samples and highly variable VDJ expression in certain tissues, and we found that both of these can lead to overly aggressive filtering by the default `cellranger` VDJ annotation and cell-calling algorithm, which is designed with a very low tolerance for doublets. Since the focus of our study prioritizes VDJ detection in VDJ -only samples over the default standard of removal of all probable doublet droplets, we designed a custom pipeline to replace the default `cellranger` VDJ algorithm for contig annotation and cell calling. This allowed us to include all high-quality VDJ sequences in the analysis, while employing more stringent filtering on samples with paired gene expression data, as described below.

We used `IgBLAST v1.17.0` to annotate all assembled contigs produced by `cellranger` and further filtered the `IgBLAST` output to retain only high quality, productive VDJ transcripts. Only contigs satisfying all of the following criteria were retained:

- `v_support` is less than exp(−60),
- `j_support` is less than exp(−10),
- V gene alignment is at least 160 nucleotides long,
- J gene alignment is at least 20 nucleotides long,
- VDJ sequence is productive,
- VDJ sequence contains a CDR3,
- VDJ sequence contains no ambiguous bases.

For all purposes except for the providing evidence for the construction of the personalized V gene databases (see below), we further removed sequences that did not contain the full-length V gene (`v_germline_start<=2`). We assigned these high quality VDJ sequences to lineages in a germline V gene reference-free way by performing single linkage clustering on sequences with the same same CDR3 length, and the same V gene family, and with the clustering condition requiring that the neither the fractional Hamming distance between the CDR3 nucleotide sequences nor the fractional Levenshtein distance between the templated regions exceeds 0.15.

We then constructed a personalized germline V gene database for each donor using grmlin [49], and re-annotated the contigs by aligning them to this personalized V gene database using BLAST v2.7.1 with the following additional options:

~~~
-word_size 9 -dust no -penalty -1 -gapopen 3 -gapextend 2.
~~~

By comparing the germline allele assignments found by `grmlin` and the germline assignments obtained by aligning the V sequence to the full IMGT database, we noticed that, in certain donors, there still remained a small number of likely germline alleles that remained undetected by `grmlin`. These were V genes that were found in their unhypermutated form in several lineages and did not represent hypermutations of other V genes in the personalized V gene database. For each donor, we ranked such additional candidate germline genes by the distance to the most similar gene in the personalized database, and then by the number of lineages the unhypermutated version of the gene was associated with. Since each of these candidate germline genes can be associated with it’s closest and, by construction, more highly expressed gene in the germline database, we further calculated the ratio, *r*, of lineages associated with each of these genes. We then iteratively added these candidate germline genes to a refined germline database if all of the following conditions were satisfied:

1. the gene is supported by at least 5 lineages, OR is at least 5 mutations away from the closest germline gene in the database, and
2. the ratio, *r*, exceeds 0.3*/d*_nearest_, where *d*_nearest_ is the Levenshtein distance between these two genes.

#### 3.2 Cell calling, V-gene tree construction, and detection and removal of cross-contaminants

##### 3.2.1 Cell calling

A significant challenge of interpreting droplet-based single-cell experiment VDJ sequencing data is distinguishing between true cells and ambient transcripts. Specifically, because the generation of droplets occurs after cells have been loaded into lysis buffer, the lysis of some cells occurs *before* they have been encapsulated in their individual droplets. As a result, a large number of transcripts are released into the ambient. Distinguishing ambient VDJ transcripts from those associated with encapsulated cells becomes challenging, especially in light of the enormous variability in VDJ expression between antibody-secreting cells and naive and memory B cells, which can differ by more than two orders of magnitude (see **Section 3.5** below). As a result, in the absence of gene expression data, it is in principle very difficult to distinguish between ambient and cell-associated VDJ transcripts.

Since one of the main quantities of interest in our study are the patterns of co-variation in the presence of different VDJs in different tissues, we employ a permissive approach to calling VDJ “cells” from this data, that we make more stringent in samples for which we do have gene expression data. Our approach has been designed with both long-read sequencing data applications (e.g. via the Pacbio platform), as well as reconstructed contigs of enzymatically fragmented amplicons (which, due to throughput limitations in long-read sequencing at the time of writing, is the exclusive source we use here, reconstructed by `cellranger`, see **Section 3.1** above). We begin by verifying that all contigs are either associated with a whitelisted 10X cell barcode, or within 1 nucleotide of an single whitelisted 10X barcode. We further discard all contigs supported by only a single UMI.

We then proceeded to assign these contigs to possible cells by iterating through all droplets and deriving a limited number of consensus contigs for each immunoglobulin locus (IGH, IGK, or IGL) using the approach described in the remainder of this section. In droplets that contained a unique VDJ per locus supported by multiple UMIs, we simply retained this contig, labeling it as a potential cell. However, as anticipated in droplet-based single cell experiments, a fraction of droplets in all samples contained contigs of the same chain that had multiple distinct VDJ sequences. These could either represent multiple encapsulated cells, ambient contaminants, or uncorrected reverse transcription, PCR or sequencing errors.

To account for uncorrected errors arising in the library generation process, we first attempted to error-correct these sequences by deriving consensus sequences for each group of VDJs in the droplet with highly similar sequences. We did this by first constructing a minimum spanning tree for the VDJs associated with each immunoglobulin chain. We examined the the distribution of distances between adjacent sequences in the minimum spanning tree, and found that it typically had a very rapid decay at small distances, and a small number of edges connecting very distant VDJs. This distribution is consistent with a small number of distinct, true VDJs surrounded by rare variants arising through errors in the library generation or sequencing process. Motivated by this observation, the minimum spanning tree was then cut by removing all edges longer than 10 nucleotides. Thus, in each droplet, for each chain, we then had a collection of graphs representing connected components of VDJ sequences which likely only differed due to library generation errors. We assigned a consensus sequence to each of these connected components via parsimony: we chose the VDJ sequence within the component that would require the smallest total number of number of mutated bases to have occurred during library preparation to explain the presence of closely-related variants.

Thus, we reduced the contigs associated with each droplet to a small number of consensus VDJ sequences that either represented multiple encapsulated cells, ambient contaminants, or a combination of the two types of species. We reasoned that ambient contaminants were likely to be supported with a smaller number of unique molecules than true cell-associated VDJs. Thus, in droplets in which there were multiple VDJ of this type, we only retained VDJ supported by at least 4 UMIs and that accounted for no fewer than 10% of all the UMIs associated with that droplet. All other VDJs were designated “ambient”.

Finally, we assigned a consensus C-gene call to each non-ambient VDJ. Due to sequence similarity among the different heavy chain isotypes, we found that, in a small fraction of cases multiple IGHC genes were associated with the same VDJ sequence in a cell. In these cases, we assigned the VDJ sequence the C-gene supported with the largest number of UMIs when that number was at least twice the number of UMIs supporting the second-ranked C-gene, and otherwise labeled the isotype as ambiguous.

##### 3.2.2 Construction of V-gene trees

To construct phylogenetic trees of non-ambient templated V sequences within a lineage, we used `MUSCLE v5.1` to first construct MSAs of all unique V nucleotide sequences within the lineage. In cases when the germline version of the gene was not present in the lineage (i.e. we did not sample a naive B cell from that lineage), we also added the germline version of the most common V gene call within that lineage to the MSA, which later allowed us to root the V gene phylogenies on this sequence. We used `fasttree v2.1.11` to infer approximate maximum-likelihood phylogenies from these nucleotide MSAs, using the generalized time-reversible nucleotide evolution model and default parameters.

#### 3.3 Light chains

In principle, IGK and IGL chains can be used to improve confidence in cell and lineage calls and to enable finer resolution of the phylogenetic tree. In practice, the inclusion of light chain information for the purposes of identifying clonal recombination events does not substantially improve accuracy [50]. Moreover, given their comparatively lower diversity and higher expression compared to heavy chains, we find that associations with cells are often substantially more difficult to distinguish from cross-contaminants or ambient species. As we detail in the next sections, such distinctions are possible for heavy chains, and so we base all of our VDJ analyses exclusively on the heavy chain sequences. However, in all samples we sequenced and bioinformatically processed the light chains, since the paired heavy and light chains sequences can be used to produce monoclonal antibodies for future experiments. These contigs will be made available alongside the contigs associated with heavy chain VDJ sequences at the time of publication.

#### 3.4 Detection and removal of cross-contaminants

Since the parallel preparation of several dozen libraries always carries the risk of cross-contamination, we endeavored to remove all potential cross-contaminating reads prior to downstream analysis. To detect cross-contamination events, we exploited the enormous diversity of both heavy chain VDJ nucleotide sequences and of 10X cell barcodes. We reasoned that, though we expect that multiple samples may contain either the same VDJ molecule or the same cell barcode, it is very unlikely that a VDJ biologically shared between two samples is also associated with the same cell barcode. Specifically, given that we typically observe on the order of 100 shared VDJs even in replicates of the same tissue, and that there are about 737,280 possible cell barcodes by 10X design, the probability that there are any accidental collisions between any given pair of samples is on the order of 6 × 10*^−^*^3^. Note that this still means that rare collisions are expected to occur in donors for which we have many samples of the same tissue, but that these collisions are expected to be rare, and that their removal should not significantly affect our estimates of rates at which sharing between tissues occurs.

**Figure S6:**
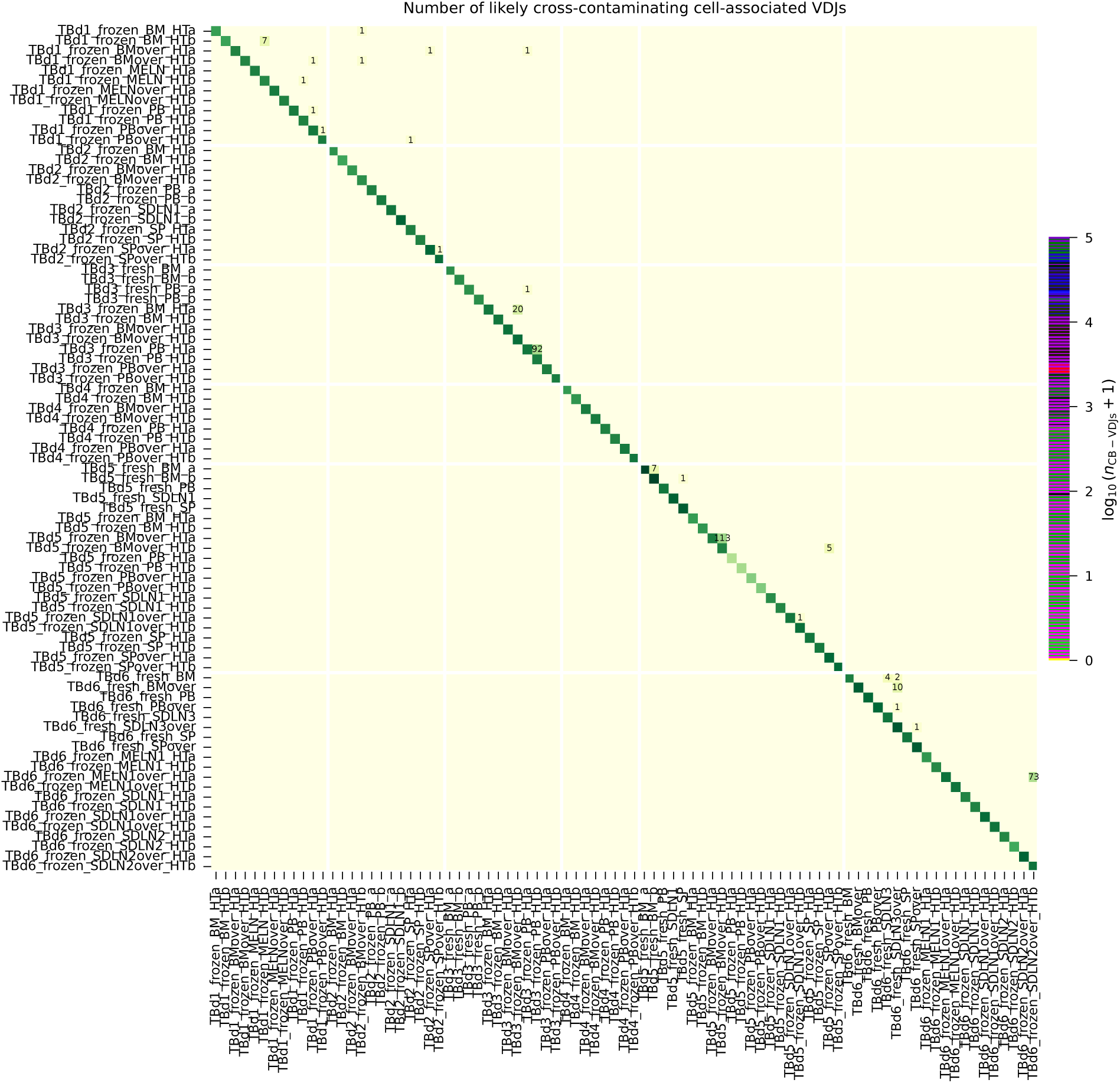
Distribution of putative cross-contaminating heavy chains. Heatmap is colored by the logarithm of the total number of likely cross-contaminating cell barcode-VDJs for each pair of samples, and by the logarithm of the total number of private cell barcode-VDJs on the diagonal. The off-diagonal elements are annotated with the number of putative cross-contaminating VDJs.

Of the 1,160,896 unique cell barcode-VDJs in our dataset, we found 348 that were shared between multiple libraries (see **Fig. S6**). In all instances, this sharing occurred between pairs of samples from the same donor or pairs of samples that that were adjacent at some point in the library preparation procedure. The shared cell barcode-VDJs were typically associated with an overall very large number of UMIs that were often concentrated in one of the samples (**Fig. S7**), indicating that they likely originate from antibody-secreting cells. This allowed us to identify one of the samples as the likely source of the contamination event. In a small fraction of cases, the shared cell-barcode VDJs were associated with a very small number of total UMIs and/or were more evenly distributed among the pair of samples (**Fig. S7**). These may represent true chance collisions of the cell barcode-VDJ pair, or may also be attributable to cross-contamination events but without a clear source sample. In this case, we removed the cell barcode-VDJ pair from further consideration. Conversely, when the shared cell barcode-VDJ was associated with a total of more than 100 UMIs, and when the fraction of UMIs in the secondary sample was smaller than 10*^−^*^2^, we removed the pair from the secondary sample and labeled the primary sample as the “source” of the contamination event.

Given that shared cell barcode-VDJs were enriched for high-UMI-count VDJs when compared to cell barcode-VDJs that were private to one of the samples (**Fig. S8**), we wanted to investigate whether there might also be a large number of undetected cross-contamination events affecting low-UMI-count cell barcode-VDJs. Specifically, since most VDJs are associated with few recovered heavy chain UMIs per cell, we were concerned that the transfer of a small number of those molecules between samples might lead to the association of that VDJ with the incorrect sample. To quantify the expected number of cross-contamination events affecting low-UMI-count VDJs, we constructed a simple model of the contamination process. Assuming that each UMI has an equivalent probability *µ* of being transferred between a pair of samples that can be involved in a cross-contamination event, the probability of a cell barcode-VDJ with *n* total UMIs being transferred between two samples (and passing filtration, which requires at least 2 UMIs to be transferred, see **Section 3.2.1**) is

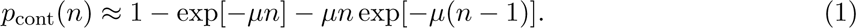

If there are *N* (*n*) cells with *n* UMIs, then we expect that the total number of contamination events is equal to

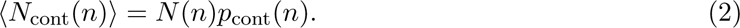

Since cross-contamination can only occur between pairs of samples that are in physical proximity, to remain conservative, we limit the estimation of *µ* and *N*_cont_(*n*) to samples in which we identified probable contamination events (pairs of shared cell barcode-VDJs). We estimate *µ* to be equal to the ratio of the sum of secondary-sample-UMIs associated with shared cell barcode-VDJs and all of the UMIs present in the pair of samples, which yields a probability of transfer per UMI of ≈ 5.9 × 10*^−^*^5^. Plugging into Equations 1 and 2, and plotting the result on **Fig. S8**, we find that our model estimates that cross-contamination events involving cells with a small number of UMIs should be extremely unlikely. Thus, we conclude that many of the low-UMI-count contamination events may indeed represent chance collisions of cell-barcodes between samples that contain identical VDJs, but that these VDJs represent such a small fraction of the sampled VDJs that they are unlikely to affect our results.

**Figure S7:**
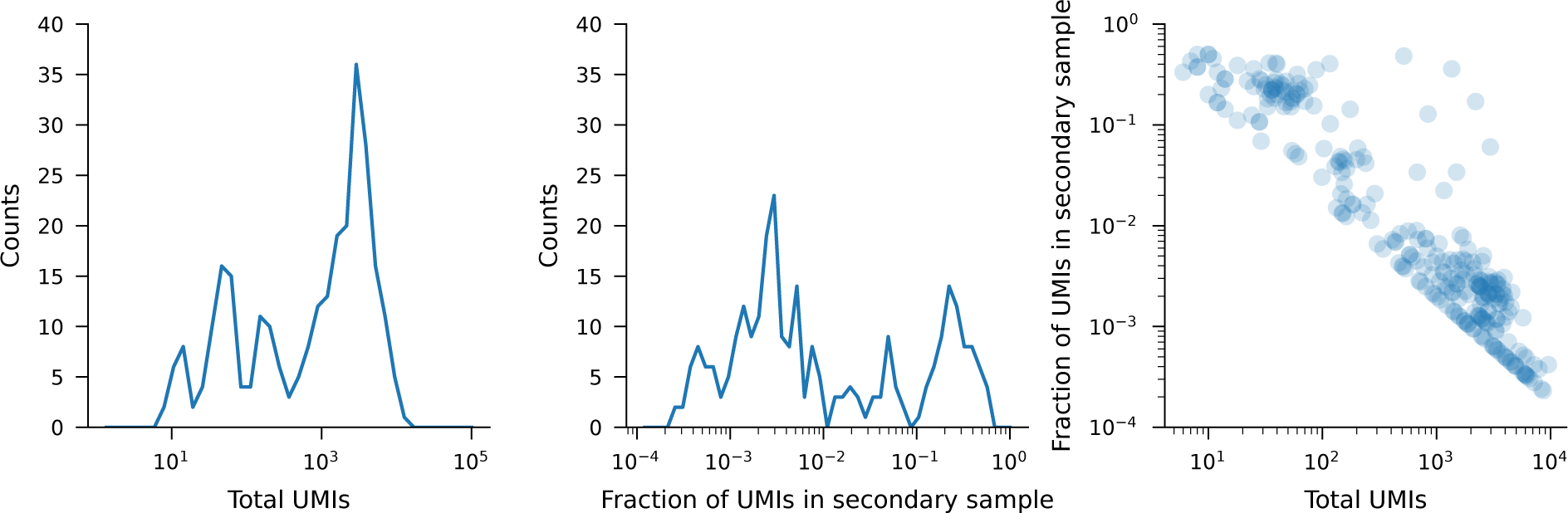
Distribution of putative cross-contaminating heavy chains among the pair of samples between which they are shared. (Left) Distribution of the total number of UMIs associated with the shared cell barcode-VDJ across the pair of samples. (Center) Distribution of the fraction of UMIs found in the sample with the smaller number of UMIs belonging to the cell barcode-VDJ pair. (Right) Scatterplot showing the two quantities for each shared cell barcode-VDJ event.

**Figure S8:**
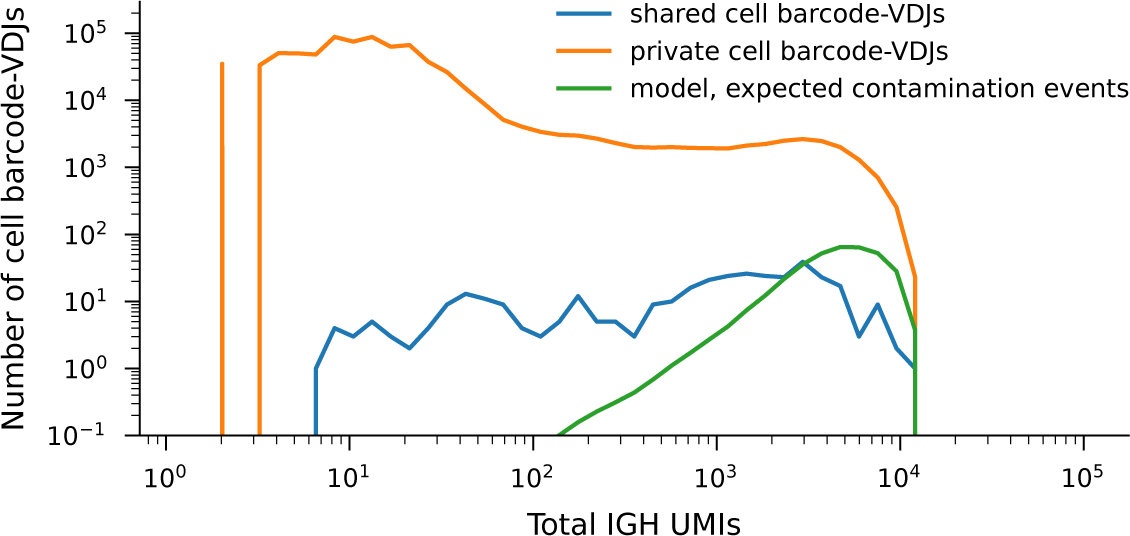
Heavy chain UMI abundance among shared cell barcode-VDJs and private cell barcode-VDJs among samples involved in any shared cell barcode-VDJ events.

#### 3.5 Transcriptome-informed analysis and annotation of ambient transcripts

##### 3.5.1 Abundance and distribution of ambient VDJ transcripts

After integrating our heavy chain VDJ transcript sequences and GEX data, we made use of the droplets associated with high-quality single non-B cells to assess the abundance of ambient VDJs in the environment. In almost all samples, a fraction of these droplets, ranging from 10*^−^*^3^ to about 0.15 contained VDJs derived from the ambient, with the majority of samples having ambient rates of a few fractions of a percent (see **Fig. S9**). The fraction of non-B cell droplets contaminated by an ambient VDJ transcript is correlated the number of high-abundance VDJ molecules in the sample (**Fig. S9**, right panel). Moreover, we find that samples thawed from storage in liquid nitrogen have no higher ambient rates than samples prepared fresh, but that there do appear to be other idiosyncratic factors that influence the rate at which the release of ambient transcripts occurs. These observations are consistent ambient antibody transcripts likely originating from antibody-secreting cells that have at least partially lysed prior to encapsulation in a droplet.

The scaling of the fraction of non-B cell droplets containing ambient VDJs with the number of high-expression VDJs also suggests that when such a lysis event occurs, it is typically limited to a small number of droplets, rather than being well-mixed in the reaction. This observation is also empirically apparent from the distribution of ambient VDJs among droplets in our data (**Fig. S10**): VDJs detected in a droplet-containing a non-B cell are typically distributed among a far smaller number of droplets than would be expected in a well-mixed model (**Fig. S10**, left panel), the majority of their UMIs are often found in a single droplet(**Fig. S10**, center panel), and more broadly, their UMIs are often highly unevenly distributed among droplets even in instances when the number of such droplets is large (**Fig. S10**, right panel). We emphasize that this spatial co-localization of ambient VDJ transcripts makes them in principle impossible to distinguish from cells in the absence of gene expression data. It also means that, in the absence of gene expression data, droplet counts are not good proxies for cell counts, and that presence of a VDJ in multiple droplets is not reliable evidence of clonal expansion.

These observations suggest that for many of these “ambient” VDJs, there might be an identifiable “source” cell, containing the majority of the VDJs and the transcriptome. Across our dataset, we find that 97% of all VDJ UMIs determined to be ambient via their association with a non-B cell transcriptome and 74% of all droplets containing an ambient VDJ can be traced back to an identifiable ASC or group of ASCs collectively carrying 78 unique VDJs. The remaining 258 ambient VDJs are typically supported by a smaller number of overall UMIs per VDJ and contain a wide diversity of VDJ sequences, without an identifiable ASC transcriptome in our dataset.

**Figure S9:**
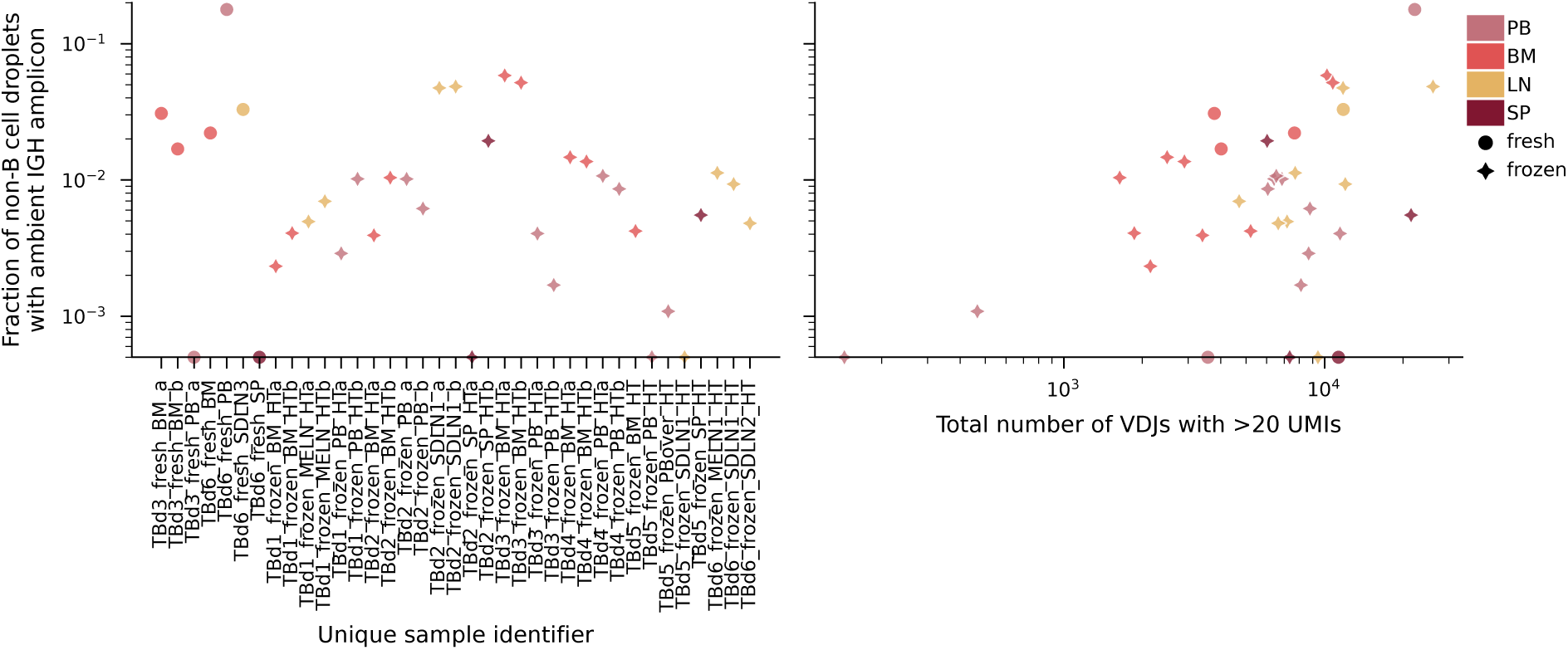
Abundance of ambient VDJs. Samples are represented by a symbol corresponding to whether they were prepared fresh or first frozen and stored in liquid nitrogen, and colored by their tissue of origin. Points on the *x*-axis represent samples in which no VDJ transcripts were found in droplets containing high quality B cells.

**Figure S10:**
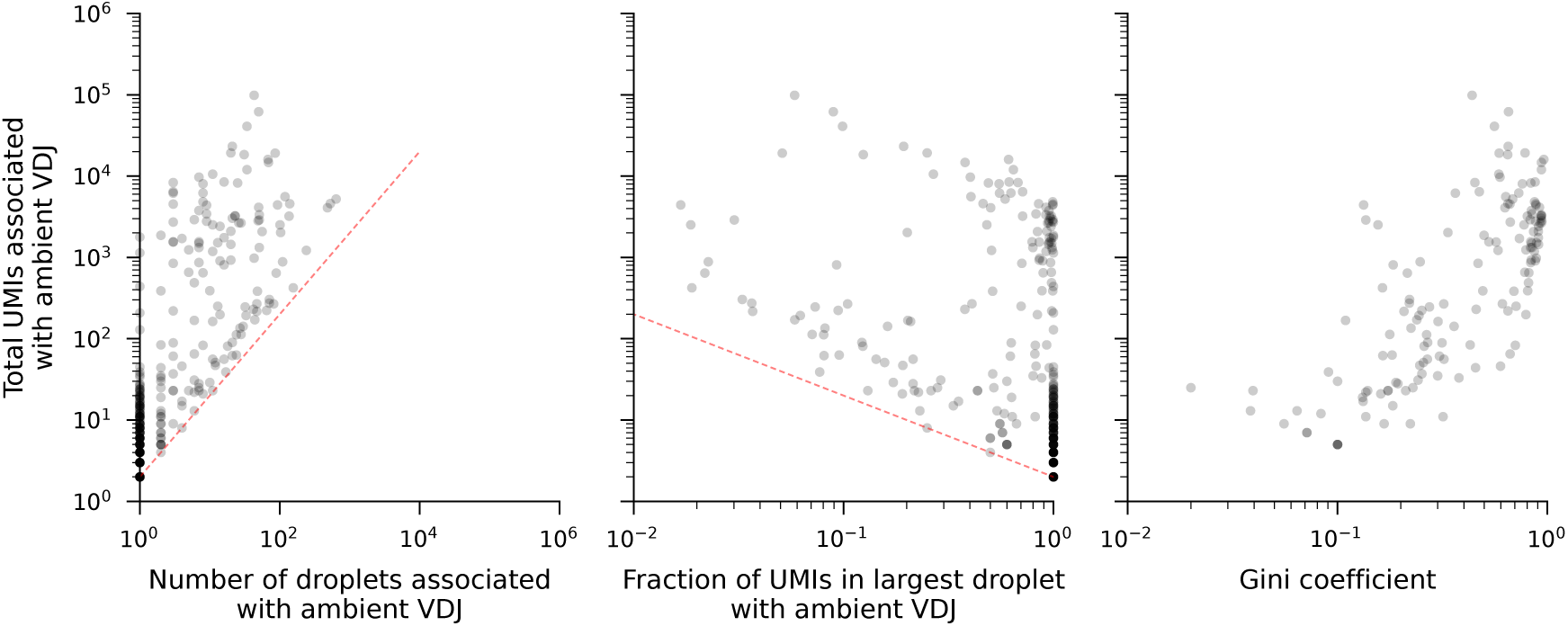
Distribution of ambient VDJs among droplets. (Left) Number of droplets and total UMIs associated with ambient VDJ. Each point represents a unique VDJ sequence associated with at least one droplet containing a high quality single non-B cell transcriptome. Red line denotes expectation under well mixed model: given the microscopic sizes of each droplet (∼ 100pL) and their enormous number in the overall volume (∼ 100*µ*L), the expected number of UMIs per droplet ranges from ∼ 10*^−^*^6^ to ∼ 10*^−^*^2^. Thus, under the well-mixed model, the probability of detecting more than the filter threshold 2 UMIs is microscopically small for the vast majority of VDJs. (Center) Fraction of UMIs in largest droplet with ambient VDJ. Dashed red line once again denotes filter threshold, expected under well-mixed model. (Right) Gini coefficient for ambient VDJs detected in more than one droplet.

Finally, we were concerned that these types of lysis events may frequently affect other ASCs in our dataset but not end up encapsulated in non-B cell droplets, making them difficult to identify as ambient. However, if we only limit ourselves to droplets that also contain a high-quality single B cell transcriptome, we find that the UMIs associated with these VDJs are found in a single droplet in over 90% of cases, and in the cases in which they are identified in multiple B-cell associated droplets, they are notably more uniformly distributed across these droplets (see **Fig. S11**. We concluded that these signatures are rather suggestive of true clonal expansion among ASCs, and do not constitute clear evidence of lysis and spillage into a number of B-cell associated droplets.

##### 3.5.2 The probability of GEX mis-identification due to the presence of ambient VDJ transcripts

Ambient VDJs detected in non-B cells can be easily removed from further analysis, but their presence in droplets associated with true B cells may lead to the identification of the VDJ sequence with the incorrect transcriptome. This is especially a concern given that many VDJ transcripts do drop out in the library preparation process (see **Fig. S12** left and discussion in **Section 1.2**).

**Figure S11:**
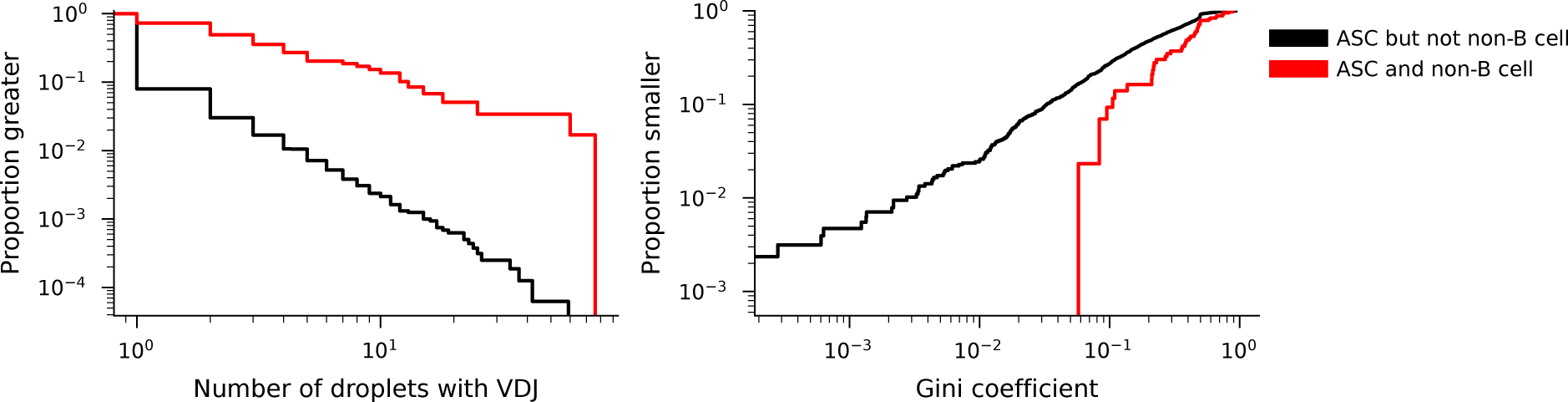
Distribution of ASC-associated VDJs among droplets. (Left) Distribution of the number of high-quality B cell droplets containing an ASC-associated VDJ for ASC-associated VDJs that either were or were not also associated with a non-B cell. (Right) Gini coefficient for ambient VDJs detected in more than one droplet.

However, we can quantify the rate at which a VDJ transcript detected in a B cell droplet is actually ambient in origin. Specifically, the empirical rate, *r*_total_, at which droplet with high-quality single B cell transcriptomes are found to also contain a VDJ transcript should be equal to the sum of the ambient contamination rates, *r*_ambient_, and the rate at which the true rate is detected in that sample, *r*_true_. Assuming that B cells are as likely as non-B cells to be associated with an ambient VDJ, the probability that the VDJ associated with the high-quality transcriptome is not ambient in origin is simply

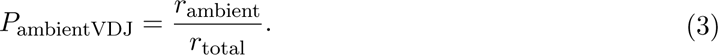

We show the distribution of these probabilities for each unique GEX sample and cell type on **Fig. S12**. Though there is significant sample-to-sample variation in the rate at which VDJs associated high-quality B cell types drop out, we find that the ambient rates are low enough that for the overwhelming majority of samples, the probability that the identified VDJ is ambient is under 5% for all B cell subsets.

Finally, we constructed an orthogonal estimate of ambient VDJs by exploiting the fact that Naive B cells should not be associated with a class-switched IGHC segment. Thus, the Naive B cell ambient contamination rate is simply equal to the ratio of the rate at which class-switched IGHC segments are found associated with Naive B cell transcriptomes and the ambient class-switched fraction, which can be estimated from the rate at which class-switched IGHC segments are found associated with non-B cell transcriptomes. These orthogonal estimates are broadly consistent with those obtained using the non-B-cell-droplet-informed (“ambient”) model (see right panel of **Fig. S12**).

##### 3.5.3 Annotation of ambient-derived VDJ transcripts

In samples for which we have GEX data, we annotated as ‘ambient’ VDJs associated with a high-quality non-B cell. In the small number of cases in which there was an identifiable ASC in the sample with the same VDJ, we annotated that cell as a ‘source’ of ambient material in cases in which the droplet in which the cell was encapsulated:

1. contained the largest number of UMIs mapping to that VDJ of all droplets that contained that VDJ,
2. contained more than 500 UMIs supporting that VDJ, and
3. accounted for more than 25% of the overall number of UMIs associated with that VDJ in that sample.

**Figure S12:**
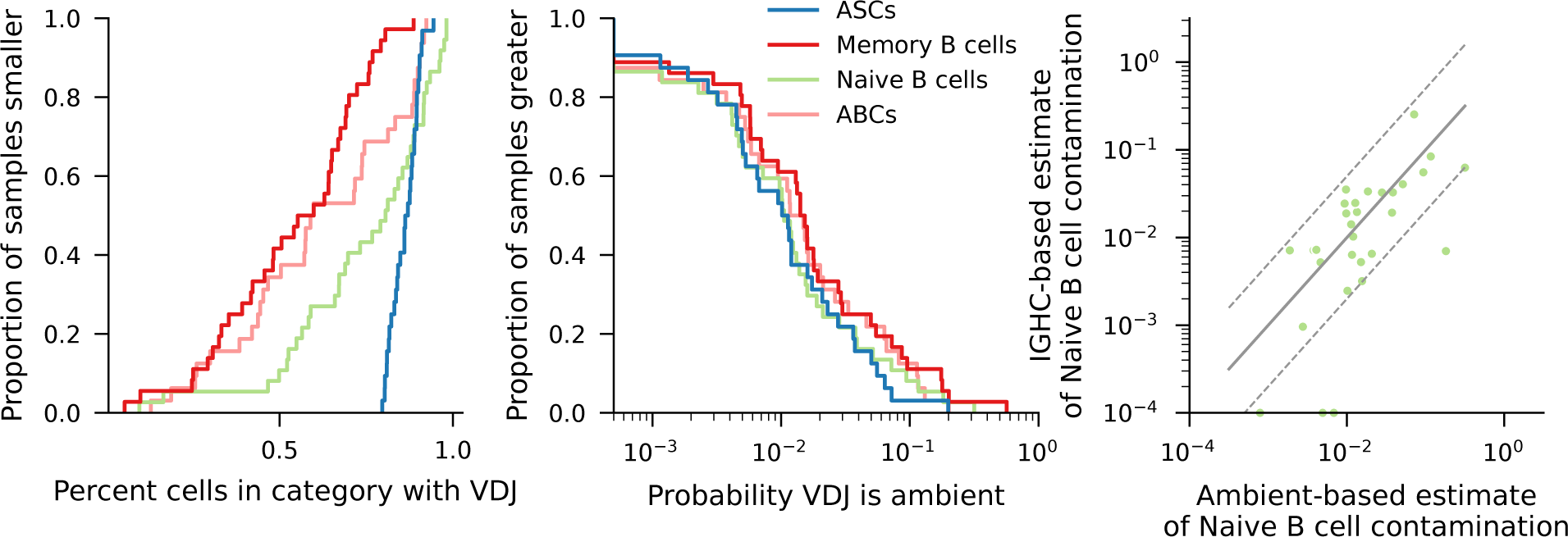
Mis-identification of ambient VDJs with GEX signatures. (Left) Distribution of rates at which a GEX profile has any identified VDJ transcripts in the dataset for each B cell subtype. (Center) The distribution of probabilities that the identified VDJ is ambient in origin for each B cell subtype. (Right) Concordance between Naive B cell droplet contamination rates from the ambient estimated based on the rates of VDJs are found associated with non-B cells (*x*-axis), or are class-switched (*y*-axis, points along the axes correspond to zeros). Each point corresponds to a unique GEX sample. The full grey line represents perfect agreement between the two estimates, and the region between the dashed grey lines corresponds to the two estimates being within a factor of 5 of each other.

### 4 Analysis of heavy chain sharing between donors

VDJ nucleotide sequence sharing between donors is remarkably rare (6 in over 600 000 unique VDJ sequences, see **Fig. ED8a**), which justifies our use of the VDJ nucleotide sequence as a unique clonal identifier within a donor. When VDJs that are shared between donors do arise, they are always seen in cells in which we do not see any hypermutations in the templated V sequence (see **Fig. ED8b**). This suggests that are associated with commonly generated CDR3s arising in independent recombination events, as opposed to convergently evolved antibodies arising in response to a common antigen. Notably, the absence of shared hypermutated VDJ nucleotide sequences suggests that even convergent recombination is far too rare an occurrence compared to the diversifying force of hypermutation to lead to the repeated generation of the identical heavy chain nucleotide sequence within the same individual. Note that this definition departs from the more commonly used definition of a “clonotype” (for heavy chains with an identical V-gene, J-gene and CDR3 amino acid sequence [51], and that it represents a far more stringent clonal identifier (**Fig. ED8c**).

To verify that our shared VDJ nucleotides sequences indeed represent convergent recombinants in naive cells, we used OLGA to calculate CDR3 generation probabilities for all CDR3s associated with unhypermutated V-gene segments[52]. Since the calculation of generation probabilities of hypermutated CDR3s also requires a model of hypermutation and selection in the germinal center reaction, which is not currently known in full, we further removed from consideration (for the purposes of this Section) all lineages in which we identified multiple unique CDR3 nucleotide sequences associated with unhypermutated VDJs. These species certainly contain hypermutated CDR3, and would lead to the erroneous inference of the generation probabilities for that CDR3, possibly skewing the entire distribution towards lower-probability CDR3’s. As we show in **Fig. ED8d**, CDR3s found in a single donor have a wide variation of generation probabilities, as inferred by OLGA. In contrast, CDR3s associated with multiple donors have exceptionally high generation probabilities, as do CDR3s associated with the 6 instances of shared VDJ nucleotide sequences between donors.

We calculated how our sharing statistics compare to those predicted by this empirical distribution of generation probabilities. To do this, we broadly followed the approach from Ref. [53], but with a slight departure in our enumeration of the number of independently generated CDR3s in our dataset. In Ref. [53], the authors, dealing with bulk TCR sequencing data, take the number of unique CDR3 amino acid sequences as a proxy for the number of unique recombination events. Here, we take the number of lineages associated with unique CDR3 nucleotide sequences and unhypermutated V genes to be the number relevant number independent rearrangements. We compare our sharing statistics to two versions of the sharing model, as in Ref. [53]. First we calculated the expected amount of sharing assuming that the probability that a productive CDR3 sequence passes negative selection at the pre-B cell stage to be order 1 (see dashed line, **Fig. ED8e**). We find that this results in comparable, but slightly lower levels of sharing than seen in our data. Therefore, we concluded that it is likely that a much smaller percentage of productive CDR3’s support high-enough quality antibodies to pass negative selection. This observation is consistent with a similar effect seen for TCRs, where only 3.7% of all productive CDR3s pass thymic selection [53]. Thus, we used our data to estimate an analogous “selection factor” for B cells, by fitting the total number of unique amino acid CDR3 sequences, conditioned on the total number of sampled independent recombination events (lineages) in each donor. We find that this factor is variable among donors, likely reflecting the paucity of data for this type of estimation, but using the mean inferred selection factor of 1.8%, we find that the empirical number of shared CDR3s lies between the two extremes of the two models we considered.

### 5 VDJ and lineage sharing analysis

#### 5.1 VDJ sharing between tissues and subanatomical regions

To construct a fair estimate of the level of VDJ sharing between all pairs of tissues and subanatomical regions, we worked to account for sample-to-sample variability in the depth of B cell sampling in our experiment by downsampling all independently generated emulsions (i.e. all independent 10X lanes) or “samples” (in the remainder of this section) to an equivalent number of 3000 unique VDJs prior to performing any comparisons. We chose this number so as to maintain the largest possible number of comparisons between pairs of tissues, and thus dropped the few extremely low-yield emulsions that contained fewer than 3000 unique VDJs from consideration in this analysis. In each sample, we obtained 100 independent samples of groups of 3000 unique VDJs, weighing each of the originally sampled VDJs equivalently (i.e. independently of the number of UMIs or droplets that the VDJ is associated with).

For each pair (*t*_1_*, t*_2_) of tissues and subanatomical regions (i.e. different individual lymph nodes), we report two quantities, the unscaled probability *p*(*t*_2_|*t*_1_) and the scaled probability *p*_normalized_(*t*_2_|*t*_1_), which we calculate as we describe below.

First, to calculate the unscaled probability that a VDJ sequence found in tissue *t*_1_ is also present in tissue *t*_2_ *p*(*t*_2_|*t*_1_), we simply calculate the average fraction of the VDJs found in tissue *t*_1_ that are also found in tissue *t*_2_, averaging over all possible pairs of *t*_2_-*t*_1_ samples derived from independent emulsions, and all replicates of each sample

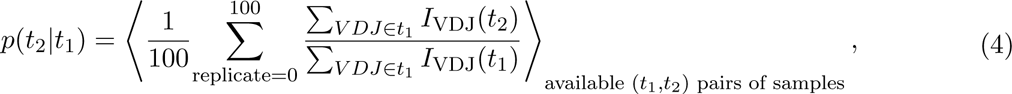

where the indicator variable *I*_VDJ_(*t*) denotes membership of the VDJ sequence in the sample of tissue *t*.

In many cases, VDJs that are truly shared between pairs of tissues may be missing simply because we are only sampling a small fraction of all the VDJs in the tissue in each sample. To account for this loss due to sampling, we scaled the raw probability *p*(*t*_2_|*t*_1_) with the probability that a VDJ found in a sample of tissue *t*_1_ is also found in an independent sample (i.e. an independent emulsion) of the same tissue (which we denote with 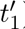). Crucially, we compute the average of the ratio of the quantities, not the ratio of the average:

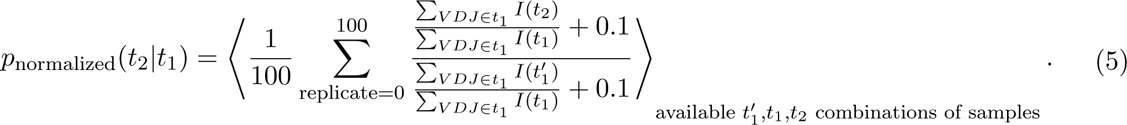

We also refer to this quantity as the“normalized” probability. We note that in the event that the distributions of the fraction of clonal cells in the tissue that carry the same VDJ are equivalent in the two tissues, and that the sampling biases during the data generation process are equivalent for the two tissues, this procedure truly scales out the sampling probability. However, even if that condition is not met, this procedure effectively conditions on the VDJ being present in a sufficiently high fraction of all cells in tissue *t*_1_ to be repeatedly sampled in independent samples of 3000 unique VDJs.

We report these quantities alongside their standard errors, computed from the variability among the different independent pairs of emulsions from tissues *t*_1_ and *t*_2_.

Finally, we also calculated the raw and scaled probabilities of discovering a VDJ in a sample of tissue *t* conditioned on it being present in a sample of two or more tissues {*t_i_*} in an analogous way:

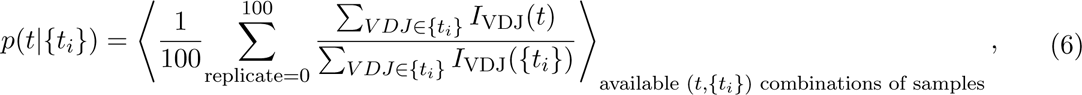

and

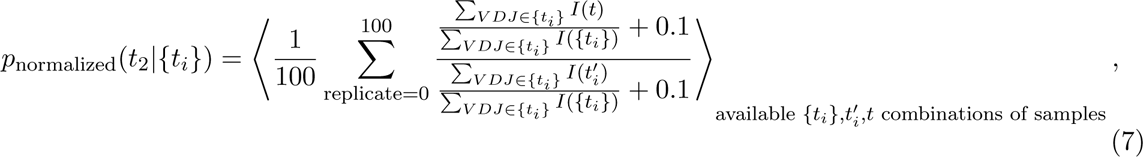

where 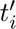 refers to an independent replicate of one of the tissues *t_i_* on which the presence of the VDJ was conditioned.

#### 5.2 Enrichment of cell types among shared VDJs

We were interested in understanding the extent to which cells in different functional states were enriched among the cells carrying shared VDJs. A VDJ was considered associated with a cell type if it was determined to be associated with a droplet containing that cell type in any of the tissues in which it was sampled.

To evaluate the enrichment in the amount of shared cells associated with a specific VDJ sequence, we constructed a simple null model that assumes that shared VDJs are not likely to be associated with any cell type in particular. This simultaneously represents a conservative model of ambient-RNA contamination of droplets of a certain cell type. In our null model, we would the expectation of the fraction of VDJs found associated with a certain cell type *c* and found in only a single tissue *t* ⟨*f*_single,c_(*t*)⟩ to be equal to the total fraction of cells of that cell type in the tissue, *f_c_*(*t*):

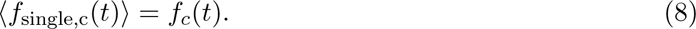

The null expectation of the fraction of VDJs found associated with a certain cell type in and found in a pair of tissues *t*_1_ and *t*_2_ is equal to

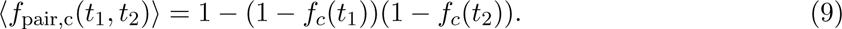

To construct the expected fraction seen in any single tissue or any pair of tissues of the same donor, we computed a weighted mean of the individual single tissue and tissue pair null expectations, where the weight was equivalent to the total number of observed cells in that tissue or tissue pair. Finally, for each quantity, we report the simple average across all donors.

#### 5.3 Gene expression associated with shared VDJs

We explored gene expression signatures in cells with shared VDJ sequences across tissues or subanatomical regions. We started by identifying genes which were differentially expressed between shared and non-shared cells, and subsequently evaluated the discriminative power of these genes. To perform differential expression analysis, we labeled each cell’s gene expression profile as “shared” between tissue pairs if its associated VDJ sequence was detected in another tissue within the same donor. We then used the Wilcoxon rank-sum test to identify genes which were differentially expressed between shared or not shared cells in each pair of tissues. In all pairs of tissues, the genes identified were indicative of an proliferative ASC phenotype, corroborating our findings in **Section 5.2**.

We were interested if there was a signature associated with the multi-tissue presence of memory B cells. To evaluate this, we performed the same differential expression analysis on only the memory B cells. For illustration, the top 12 differentially expressed genes for the Lymph Nodes and the Spleen are shown **Fig. ED10a**). Upon considering other pairs of tissues, we noticed that many of them had similar sets of differentially expressed genes. Thus, we used the 45 most differentially expressed genes between shared memory B cells in the Lymph Nodes and Spleen as the features to train a logistic regression classifier for sharing across all tissue pairs. Before training, we downsampled the data to remove the class imbalance in the sharing variable, because a large majority of cells were not shared. We found that this classifier could be used to enrich for shared memory B cells across any pair of tissues, suggesting that there is a common gene expression profile associated with memory B cell presence in multiple tissues (**Fig. ED10c**), though it is difficult to report a measure of the false-negative and false positive rates associated with it, because many truly shared VDJs will not have a ‘shared’ label in the context of limited sampling. We report the inferred scale parameters (‘feature importances’) for each of the features included in the classifier in **Fig. ED10b**.

#### 5.4 Lineage sharing between tissues

We quantified the level of lineage sharing between tissues using an entirely analogous approach to the one we used to quantify VDJ sharing described in **Section 5.1**, with two minor differences:

1. all samples were downsampled to 5000 unique VDJs prior to estimating all quantities, and samples in which we recovered fewer than 5000 unique VDJs were dropped from the analysis. The reported quantities represent averages of all possible pairs of samples informing the comparison between two tissues, and were computed based on a single random subsample, and
2. we computed normalized probabilities of sharing of lineages conditioned on their binned frequency in one of the tissues.

We emphasize that, with the exception of TBd1, in whose peripheral blood and bone marrow we find evidence of a substantial excess of high-frequency lineages, lineage size distributions collapse after subsampling, justifying the scaling procedure used above to account for subsampling.

### 6 A model of B cell differentiation during affinity maturation

The uniform distribution of hypermutation levels among related cells in distinct differentiated states suggests that B cell differentiation biases do not vary as the germinal center reaction proceeds: as germinal center (GC) B cells undergo rounds of selection during affinity maturation, their relative probabilities of differentiating into a Memory B cell or an ASC remain constant. These data contrast with a view in which there is an explicit temporal variable that governs the probability of differentiating into a memory B cell (reviewed in Ref. [15]), and favors a model in which the affinity of a B cell for the antigen is the primary determinant of GC B cell fates. However, to be taken seriously, this model of the GC reaction must also be capable of explaining the higher average hypermutation levels in ASC’s compared to Memory B cells, as well as their overall lower abundance. In this section, we propose a simple model of the B cell fate choice in the GC reaction that is capable of reproducing all of these features.

In the simplest model of the GC reaction that is consistent with the observation of uniform distributions of hypermutation levels, the GC reaction proceeds as a memory-less process in which all cell fate decisions are constant in time. To elaborate, in each round of selection in the light zone of the germinal center, GC B cells may differentiate and exit the GC as Memory B cells or ASCs, reenter the dark zone for continued hypermutation, or fail to receive signals for continued survival and proliferation and undergo apoptosis. Crucially, the probability of each of these events does not change as the GC reaction proceeds. We illustrate this process schematically in **Fig. S13**.

Though the full specification of this model requires us to also specify a model of proliferative bursts in the dark zone of the GC, which is beyond the scope of this work, we can still make conceptual progress by making several crude approximations. Assuming that a B cell accumulates on average *V* hypermutations in the templated portion of the IGHV gene in each GC cycle, that the GC itself neither grows or shrinks substantially after the GC reaction is established, and that different cell types do not undergo substantially different proliferative bursts, we can approximate the probability that a hypermutated B cell in state *i* has *m* or more hypermutations after a single germinal center reaction as

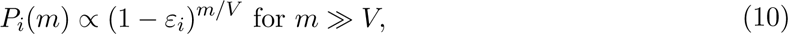

where *ε_i_* is the effective exit rate from the germinal center in state *i*. The relationship between this quantity and the parameters *δ* and the differentiation rates *θ_k_* depends on whether or not all cell fates are in principle accessible on all mutational paths and in all germinal centers, in some statistical sense. If that is the case, then for all exit fates *ε_i_* = *δ* + Σ*_k_ θ_k_*, where the summation is over all possible fates. In the opposite extreme, in which the paths that lead to an exit in state *i* make other events inaccessible, *ε_i_* should reach its lower bound for that event, *θ_i_*, the differentiation rate to state *i*. This could either arise as a result of a form of statistical contingency in the binding landscape of the antibody, an intrinsic property of the initial cell state, or due to a contingency in the form of differentiation signals available to that cell in the germinal center (e.g. the availability of certain cytokines, or the specificity of the available T cells in the germinal center). These different processes are not fully identifiable from observational data alone, but we can nonetheless make progress by inferring the effective exit rates *ε_i_*(*m*) and observing how they change over time and how they vary between different exit fates.

**Figure S13:**
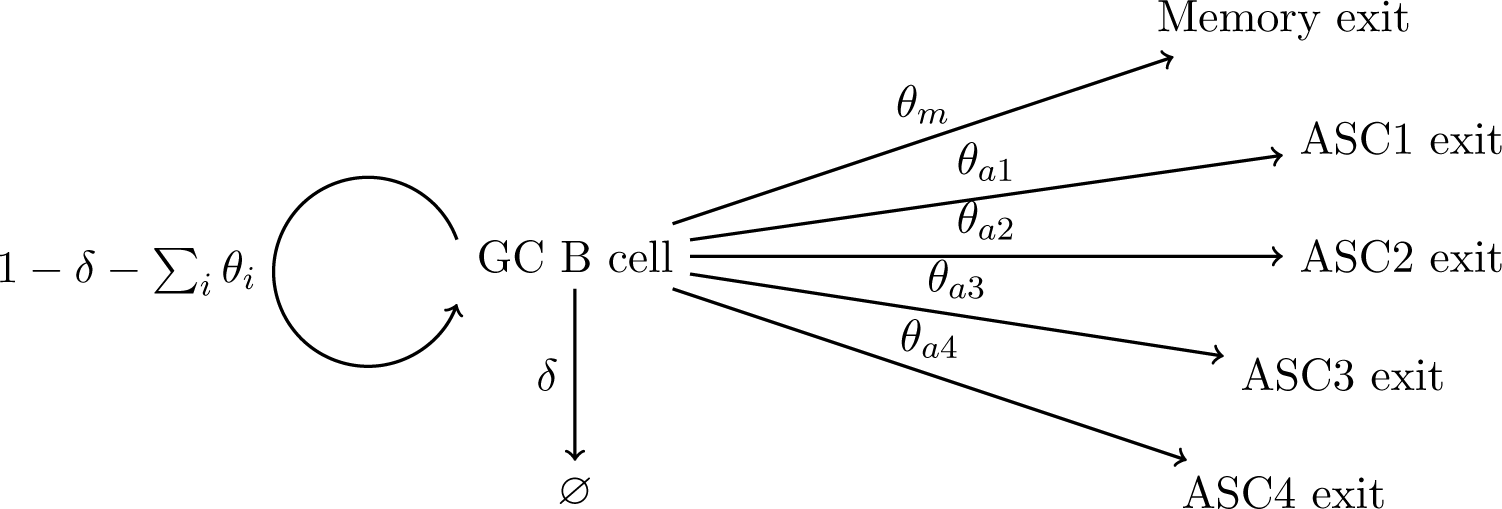
Schematic illustrating a model of a memoryless differentiation process within an active germinal center. The per-LZ/DZ cycle probabilities of each of the events are denoted by the symbols accompanying each arrow.

**Figure S14:**
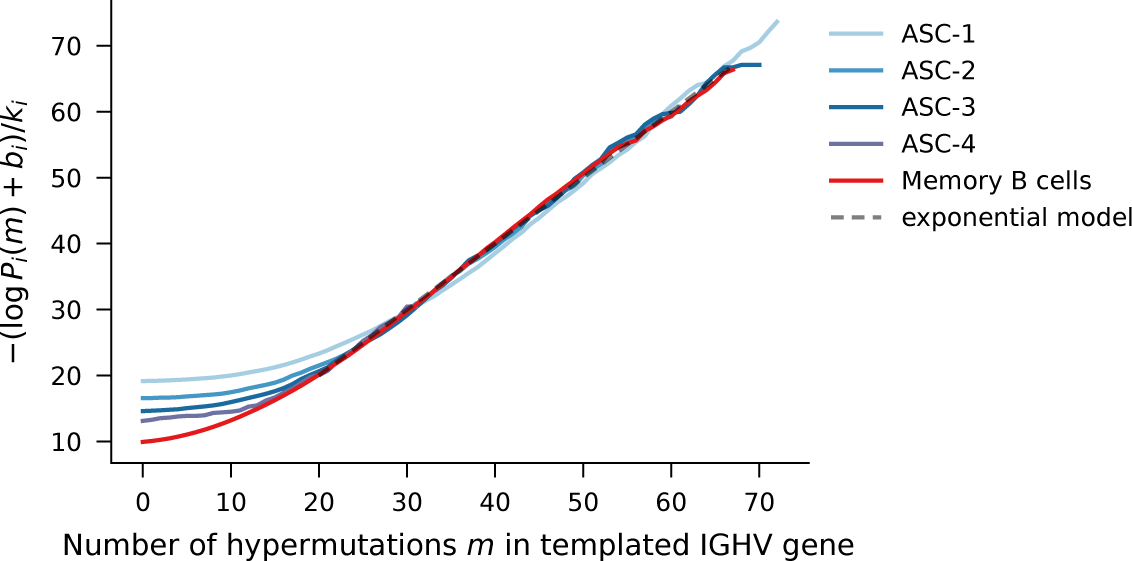
Scaled logarithm of the complementary CDF. Theoretical expectation based on model in **Fig. S13** is indicated by the dashed grey line. Parameters *k_i_* and *b_i_* for each cell type were inferred inferred by least-squares linear regression of the logarithm of the complementary CDF for *m* ≥ 20. Portions of the CDF supported by fewer than 20 cells have been truncated.

If the effective exit rates are constant over time (the simplest verision of this model), at sufficiently large *m* (such that we can be confident that the cell has undergone many GC cycles), the negative logarithm of the complementary cumulative density function (CDF)) is expected to increase linearly with the number of hypermutations *m*

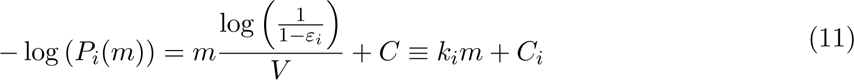

with slope *k_i_*that is related to the exit rate *ε_i_* according to

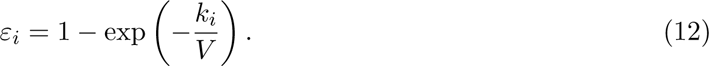

We compare the predictions of this constant rate model with the distributions of hypermutation levels in our data in **Fig. S14**. The inferred slopes *k_i_* and the corresponding per-cycle exit rates *ε_i_* obtained from *k_i_*assuming that cells undergo on average 1 division per cycle (*V* ≈ 0.3, [54, 55]) are shown in **Table S4**. Notably, all of the inferred slopes, and therefore also the exit rates, *ε_i_*, are very similar, but not identical. This suggests that not all paths and germinal center reactions are fully exchangeable in some statistical sense. In particular, we cannot exclude that parts of the population of cells destined to become memory B cells has a somewhat different set of possible internal states that are inaccessible to cells destined to become ASCs. In contrast, the differences in exit rates between the ASC subtypes are smaller, and thus it appears that cells differentiating into ASCs may potentially have simultaneous access to all different ASC fates.

On examining the shapes of the theoretical and empirical distributions of mutational levels, we find agreement over several orders of magnitude in the case of Memory B cells and ASCs that have accumulated at least 10 hypermutations. However, we also find a modest reduction in the number of weakly hypermutated memory B cells and ASCs compared to the expectations of this model. The rate at which early ASCs exit from the germinal center is similar for ASCs of different types and lower than that for memory B cells by up to a factor of 2, but becomes indistinguishable for the differentiated B cell types after they have accumulated about 10 V gene hypermutations (corresponding to about 3% divergence from germline). We note that in lineages that are sufficiently large for us to be able to make comparisons between related cells, the relative fraction of cells with fewer than 10 V gene hypermutations is very small. Specifically, among lineages for which more than 3 unique VDJs have been sampled in our dataset fewer than 20% of all VDJ sequences have fewer than 10 hypermutations, and so the majority of the cells in these lineages have exited the germinal center at a time when our data indicate differentiated cells exit it with a constant probability at every GC cycle (**Fig. S15**).

We propose two potential hypotheses for the absence of weakly hypermutated differentiated B cells. First, it is possible that cells destined to become differentiated B cells undergo a larger proliferative burst in the dark zone of the germinal center, acquiring potentially a larger number of hypermutations in each GC cycle (corresponding to a larger *V* parameter in our model, since proliferation and hypermutation are coupled). This alone would make the exit of a differentiated B cell at a small number of hypermutations less probable, since return to the light zone is thought to be necessary for differentiation signals. Moreover, the proliferative burst size and therefore the number of hypermutations acquired in each cycle may vary between the different exit states, and may be lower for memory B cells than for ASCs. However, as long as the distribution of these burst sizes for cells of the same type is not too broad, we expect this model to yield a distribution indistinguishable from the one considered above at large *m*.

Second, it remains possible that at very early stages of the GC reaction, there remains explicit time-dependence in the rate at which GC cells differentiate into memory B cells and ASCs, as proposed before by other authors (see Ref. [15]). If this is the case, our data suggests that this time-dependence is limited to very early stages of the GC reaction, and that 80% of all differentiated cells exit the GC at times when this effect is negligible.

**Table S4:** Inferred per-cycle exit rates for the model implied in Fig. S13.

| Exit state | Inferred slope, $k_i$ | Per-cycle exit rate $\varepsilon_i$ |
| --- | --- | --- |
| ASC-1 | 0.10 | 0.030 |
| ASC-2 | 0.10 | 0.029 |
| ASC-3 | 0.11 | 0.033 |
| ASC-4 | 0.10 | 0.029 |
| Memory B cells | 0.14 | 0.041 |

**Figure S15:**
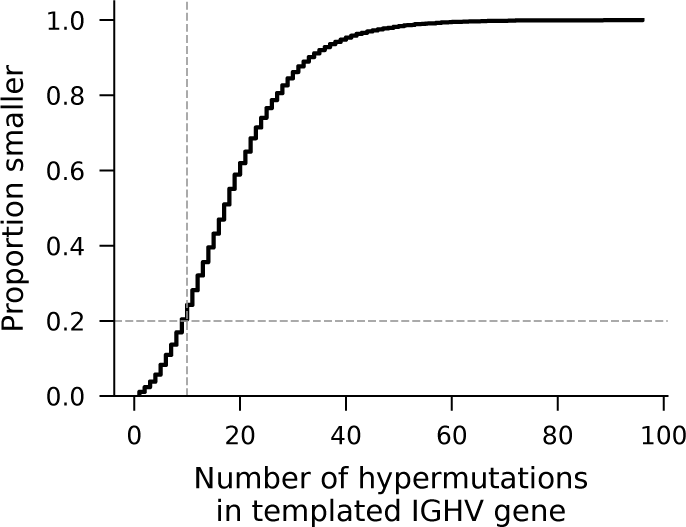
Distribution of hypermutation numbers in all cells found in lineages with at least 3 sampled unique VDJs. Dotted lines indicate point at which the ratio of Memory B cell GC exit rates and ASC GC exit rates appears to stabilize.

